# A single short reprogramming early in life improves fitness and increases lifespan in old age

**DOI:** 10.1101/2021.05.13.443979

**Authors:** Quentin Alle, Enora Le Borgne, Paul Bensadoun, Camille Lemey, Nelly Béchir, Mélissa Gabanou, Fanny Estermann, Christelle Bertrand-Gaday, Laurence Pessemesse, Karine Toupet, Jérôme Vialaret, Christophe Hirtz, Danièle Noël, Christian Jorgensen, François Casas, Ollivier Milhavet, Jean-Marc Lemaitre

## Abstract

Forced and maintained expression of four transcription factors OCT4, SOX2, KLF4 and c-MYC (OSKM), can reprogram somatic cells into induced Pluripotent Stem Cells (iPSCs) and a limited OSKM induction is able to rejuvenate the cell physiology without changing the cell identity. We therefore sought to determine if a burst of OSKM might improve tissue fitness and delay age-related pathologies in a whole animal. For this, we used a sensitive model of heterozygous premature aging mice carrying just one mutated Lamin A allele producing progerin. We briefly treated two months-young heterozygotes mice with OSKM and monitored their natural age-related deterioration by various health parameters. Surprisingly, a single two and a half weeks reprogramming was sufficient to improve body composition and functional capacities, over the entire lifespan. Mice treated early in life had improved tissue structures in bone, lung, spleen, kidney and skin, with an increased lifespan of 15%, associated to a differential DNA methylation signature. Altogether, our results indicate that a single short reprogramming early in life might initiate and propagate an epigenetically related rejuvenated cell physiology, to promote a healthy lifespan.

**One Sentence summary:** A single short reprogramming early in life rejuvenates cell physiology, improves body composition, tissue fitness and increases lifespan in elderly.

## Introduction

Over the course of a human lifetime, the entry into old-age brings increased likelihood of contracting age-related diseases. The aging demographic makes this issue a central scientific concern in medicine. Aging is a complex process often punctuated by the appearance of age-related pathologies and a decrease of cell and tissue regenerative capacity. It intensifies cell and tissue vulnerability and deterioration and increases the risk of developing diseases like cancer, cardiovascular disorders, diabetes, atherosclerosis, age-related macular degeneration or neurodegeneration and ultimately precipitating death (1, 2). The mechanisms causing aging are still poorly understood, making it difficult to develop prophylactic strategies to increase healthy lifespan. There are numerous molecular and cellular hallmarks of the aging process, including, cellular senescence, genomic instability, deregulated autophagy, mitochondrial dysfunction, telomere shortening, oxidative stress, systemic inflammation, metabolism dysfunctions, epigenetic alterations and stem cell exhaustion (2). Consistently, the capacity of youthful tissue to self-renew, through the action of stem cells is impaired by senescence. Among the described hallmarks, DNA methylation was proposed to be pertinent to evaluate the physiological age of individuals, since the deviation of predicted and chronological age correlates with all-cause mortality in human (3, 4) and was described to be affected by genetic, dietary, or pharmacological interventions (5-7). Although many of these hallmarks have been extensively described and studied, few of them have been translated into effective therapies, with the notable exception of the removal of senescent cells, which has led to the development of senolytic drugs in humans (8).

In 2006, it was shown that mouse somatic cells can be converted into pluripotent cells (iPSCs) by inducing the expression of four transcription factors: OCT4, SOX2, KLF4 and c-MYC (OSKM) (9). This process of cellular reprogramming induces a global remodeling of epigenetic landscape to revert cell identity to a pluripotent embryonic-like state. Exploiting cell reprogramming offers an alternative route for cell therapy to restore organ and tissue function. Somatic cells can be reprogrammed into iPSCs, then modified or corrected *in vitro* before being re-differentiated into cells, tissues or organs for replacement in the donor or an immune-compatible patient (10). Recently, a new reprogramming method was developed using a transient expression of a nuclear reprogramming factors to promote amelioration of aging hallmarks human cells (11).

Previous experiments using a reprogrammable mouse model demonstrated that a cyclic induction of OSKM two days a week, over the entire extremely short lifetime of a homozygous accelerated aging mouse model, increased longevity, through a potential chronically unstable epigenetic remodeling (12). These mice have a mutated *Lmna* gene that produces high level of the natural aging protein progerin (13).

In this study, we investigate for the unexplored of a single short period of *in vivo* OSKM induction as pre-clinical proof of principle for a potential usage in clinic to prevent aging defects. We focused on heterozygous animals, which have moderate lifespan and levels of progerin (14), as these heterozygotes might be extremely sensitive to anti-aging therapies. As a short OSKM induction, was described to ameliorate immediate tissue regeneration after experimentally induced tissues injuries, we wondered whether a short period of OSKM genes induction might improve lifespan and tissues aging of heterozygotes mice. Surprisingly, we found that many health measures, and longevity itself, were ameliorated in elderly, by a single two and a half weeks treatment on two months old mice, associated to a differential DNA methylation signature, suggesting that a “memorized effect” initiated by our short induction protocol early in life might be involved in a more juvenile physiology.

## Results

### A single early short transient reprogramming increases late age lifespan

We considered a specific mouse model (Lmna^G609G/+^) recapitulating the human phenotype of HGPS mimicking accelerated physiological aging due to a point mutation (G609G) in the *Lmna* gene (14). As a consequence, the protein Progerin, a truncated form of Lamin A responsible for human HGPS (15), accumulates and mice are short-lived. Homozygous mice have an average lifespan of 15 weeks while heterozygous mice live around 35 weeks. In both cases, due to accelerated aging mice also present accelerated onset of several phenotypic alterations related to aging notably including weight loss and lordosis. Lamin-compromised mice also have damages to skin, liver, kidney, bone and heart (14). We crossed these mice with a homozygous transgenic murine model (R26^rtTA/rtTA^;Col1a1^4F2A/4F2A^) allowing, the controlled induction of the expression of OSKM factors through an rtTA trans-activator, thus reprogramming all the animal’s cells (16). Addition of doxycycline (DOX) in the drinking water of the triple heterozygous transgenic mice (R26^rtTA/+^;Col1a1^4F2A/+^;Lmna^G609G/+^) allows the controlled expression of the four reprogramming factors to establish reprogramming protocols in mice mimicking accelerated physiological aging.

Firstly, we sought to revisit the previously published OSKM treatment, which was to administrate 1 mg/ml DOX, two days a week, throughout life on our heterozygous progeric model (12). Thus, we tested a simplified protocol, where a reduced concentration was added continuously throughout life, and used the previously published chronic induction protocol as a reference. Strikingly, this simplified protocol gave near identical improvement to heterozygous lifespan than the chronic one, with a high increase in median age of death, from 42.6 to 55.6 weeks, meaning that lower doses of doxycycline could be effective (Figure 1A, Supplementary Figure 1A). There was also a significant reduction in age-related weight loss, when compared to non-induced control animals (Supplementary Figure 2A).

**Figure 1:**
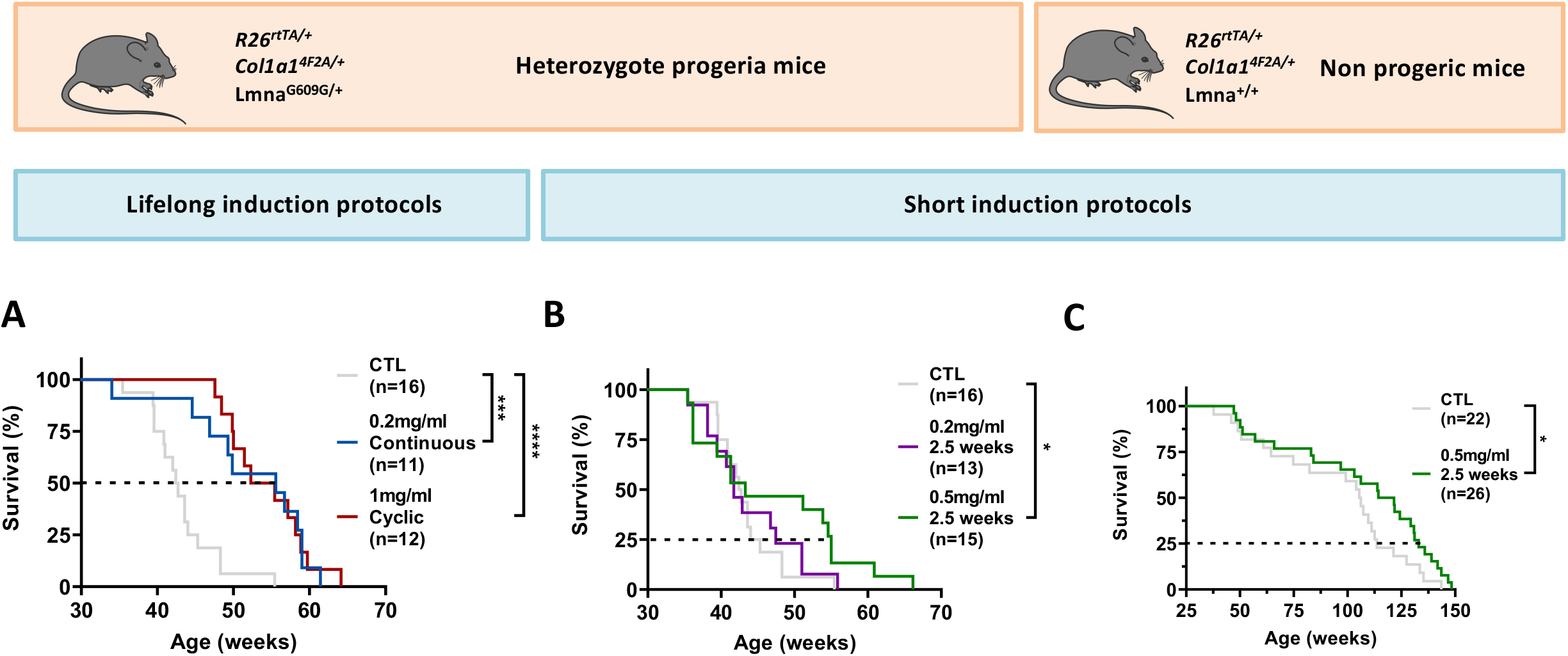
Evaluation of the different reprogramming induction regimens on lifespan. **(A)** Long-term OSKM induction protocols were performed on progeric R26^rtTA/+^;Col1a1^4F2A/+^;Lmna^G609G/+^ mice by administrating 0.2 mg/ml doxycycline in the drinking water either continuously (blue curve) or a chronically 2 days a week (red curve), started at 2 months old and maintained over the entire life. Survival curves of long-term doxycycline-treated mice after induction compared to untreated mice (grey curve) with the same genotype are presented. Statistical analysis of curves was performed at the corresponding indicated percent survival. **(B)** Short-term OSKM induction protocols were performed on progeric R26^rtTA/+^;Col1a1^4F2A/+^;Lmna^G609G/+^ mice, by administrating doxycycline in the drinking water for 2.5 weeks either at 0.2 mg/ml (purple curve) or at 0.5 mg/ml (green curve). Inductions start at 2 months old. Survival curves of short-term doxycycline treated mice after induction compared to untreated mice with the same genotype are presented. Statistical analysis of curves was performed at the corresponding indicated percent survival. **(C)** Short-term OSKM induction protocol was performed by adding doxycycline in the drinking water for a single period of 2.5 weeks at 0.5 mg/ml (green curve) on non-progeric R26^rtTA/+^;Col1a1^4F2A/+^;Lmna^+/+^ mice. Survival curves of short-term doxycycline treated mice after induction compared to untreated mice (grey curve) with the same genotype are presented. Statistical analysis of curves was performed at the corresponding indicated percent survival. * p=0.0113, *** p=0.0003, **** p<0.0001; according to log-rank (Mantel-Cox) test.

Then, we further investigated whether an extremely simplified protocol of a single period of treatment might cause a permanent improvement in vitality, as it might be straightforward for clinical translation. We thus induced reprogramming factors with of 0.2 mg/ml and 0.5 mg/ml DOX for just a single two and a half weeks treatment at two months of age (equivalent to adolescence in mice). There was a very minor but non-significant effect at 0.2 mg/ml. However, the short more concentrated treatment noticeably improved lifespan in those mice surviving to middle age (Figure 1B).

Indeed, while no difference was noticeable in median age of death, the 0.5 mg/ml doxycycline treatment remarkably increased the age of death for the third quartile from 44.3 weeks to 54.8 weeks for treated animals compared to controls (Figure 1B, Supplementary Figure 1B). Surprisingly, this protocol also increased the maximum lifespan in the group to 66.1 weeks, which is 11 weeks longer than the longest-lived animals (Figure 1B, Supplementary Figure 1B). Consistently, there was a significant reduction in age-related weight loss, when compared to non-induced control animals (Supplementary Figure 2B). We finally develop this last protocol on non-progeric mice and we observed a similar but smaller effect on longevity, consistent with a lower sensitivity of healthy animals. Wild type mice that survive to middle age (the third quartile) lived on average 113 weeks instead of 131 weeks in the treated group (Figure 1C).

Collectively, these results demonstrate for the first time that a single short transient expression of reprogramming factors *in vivo* can increase lifespan of progeric and non-progeric mice.

### *In vitro* transient reprogramming rejuvenates cell physiology

As the DOX inducible OSKM cassette is present in all cells of our mice models, we decided to further analyze the impact of our short induction protocol, in a simple *in vitro* cellular model. OSKM induction from just one allele does not force derivation of iPSCs and do not form teratomas in mice as previously demonstrated by others in a different reprogramming model (17).

To further investigate for the consequences of OSKM, we firstly evaluated the immediate impact of an OSKM induction in cell physiology. We collected adult skin fibroblasts from reprogrammable non-progeric control mice (R26^rtTA/+^;Col1a1^4F2A^;Lmna^+/+^).

Interestingly, induction of OSKM decreased DNA damages, as shown by γ-H2AX level, as well as a lower expression of FOXO3, that has been shown to be activated in stress response situations (Figure 2A) (18). A significant increase in cell proliferation was also detected without any increase in cell death (data not shown). Senescence also decreased as demonstrated by reduced SA-β-Galactosidase staining (Figure 2B). In parallel to this observed amelioration on cell physiology, we observed an increased autophagic flux which is known to recycle altered cell components like carbonylated proteins or damaged mitochondria (Figure 2C) (2, 19). Unexpectedly, we measured a two-fold increase of the mean number of mitochondria per cell, which could be not attributed to an immediate boosting effect on metabolism since glycolysis and Krebs cycle metabolites level remained constant (Figure 2D). We speculated that this might improve cell metabolism to respond to future energy demands in stress situations. Overall, our results are consistent with an immediate rejuvenation effect of OSKM induction on cell physiology.

**Figure 2:**
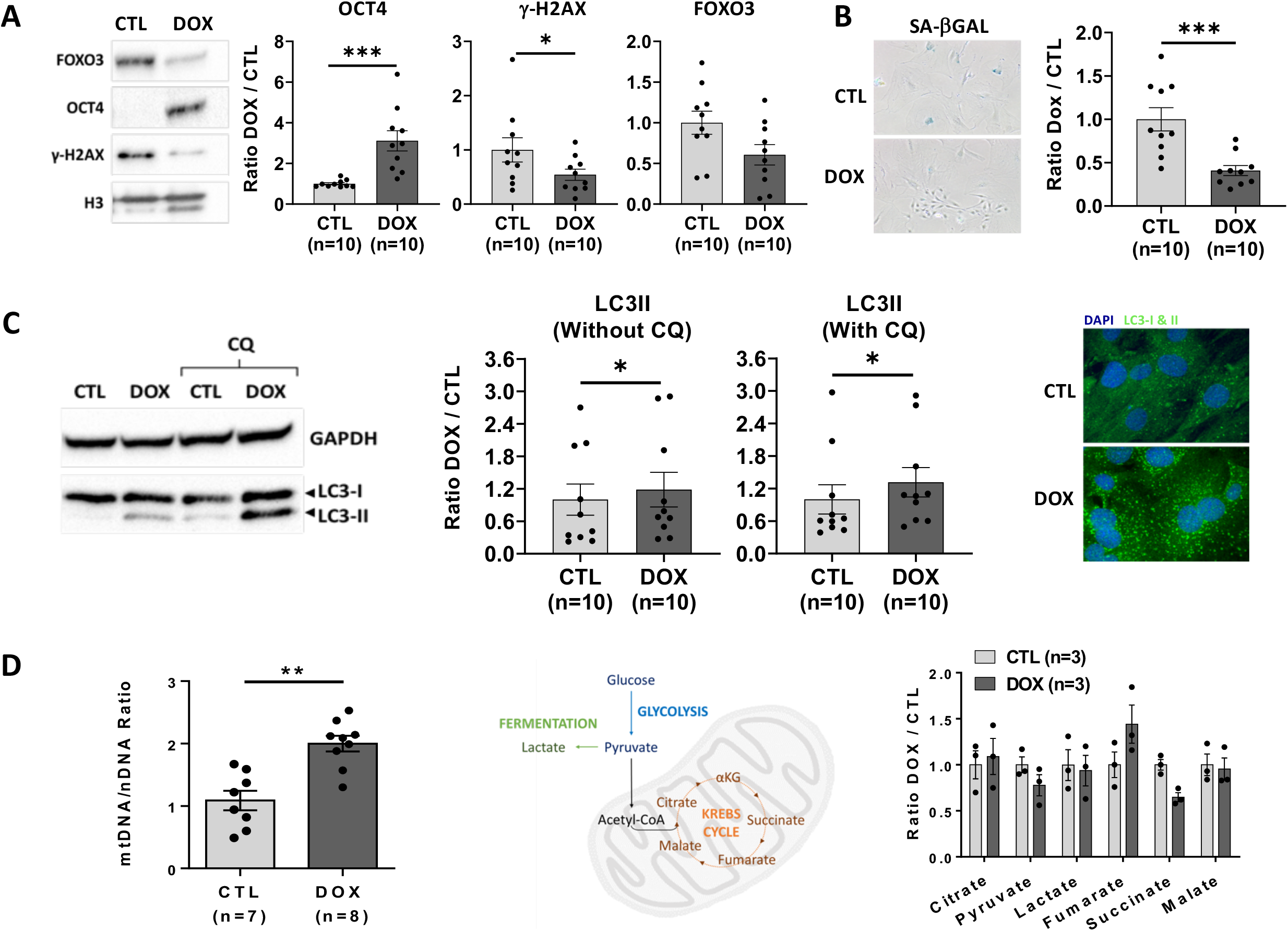
Rejuvenating effect of a short OSKM burst on cell physiology. Cell physiology parameters were analyzed in skin fibroblasts. **(A)** Damage and stress response visualized by *γ*-H2AX and FOXO3 protein levels in treated (DOX) compared to non-treated cells (CTL). Protein levels were detected by western blot and signals were quantified after normalization by H3 signal. According to the number of samples, cell lysates were prepared all together, run on different gels and then further processed identically before quantitative comparison. **(B)** Senescence evaluated by the percentage SA-βGAL staining positive cells in treated (DOX) compared to non-treated cells (CTL) **(C)** Evaluation of the autophagic flux by the level of LC3II in presence or absence of chloroquine (CQ) in DOX-treated compared to non-treated cells (CTL), quantified by immunoblot. Immunocytochemistry depicting the increased level of LC3I&II in treated (DOX) compared to non-treated cells (CTL). According to the number of samples, cell lysates were prepared all together, run on different gels and then further processed identically before quantitative comparison. **(D)** Quantification by qPCR of the mtDNA copy number per cell in OSKM induced skin fibroblasts and quantification of the level of some metabolites related to energy production, by metabolomics.

### *In vitro* transient reprogramming induces gene pathways targeting specific aging diseases

To further explore for the causes of the increased longevity in progeric and non-progeric mice, we collected adult skin fibroblasts from reprogrammable heterozygous progeria mice (R26^rtTA/+^;Col1a1^4F2A/+^;Lmna^G609G/+^) and non-progeric control mice (R26^rtTA/+^;Col1a1^4F2A^;Lmna^+/+^) and induced the four reprogramming factors OSKM by doxycycline. Gene expression profiles of OSKM-induced skin fibroblasts from both genotypes and sexes were then analyzed by RNAseq (Figure 3A, Supplementary Figure 3). 640 and 533 genes were differentially expressed in both sexes in heterozygous and wild type fibroblasts respectively by inducing OSKM (Figure 3B). 395 of these genes were differentially expressed in both genotypes and sexes (Figure 3B). Thus, independently of genotypes and sexes, cells displayed a distinct pattern of gene expression after treatment with doxycycline (Figure 3C).

**Figure 3:**
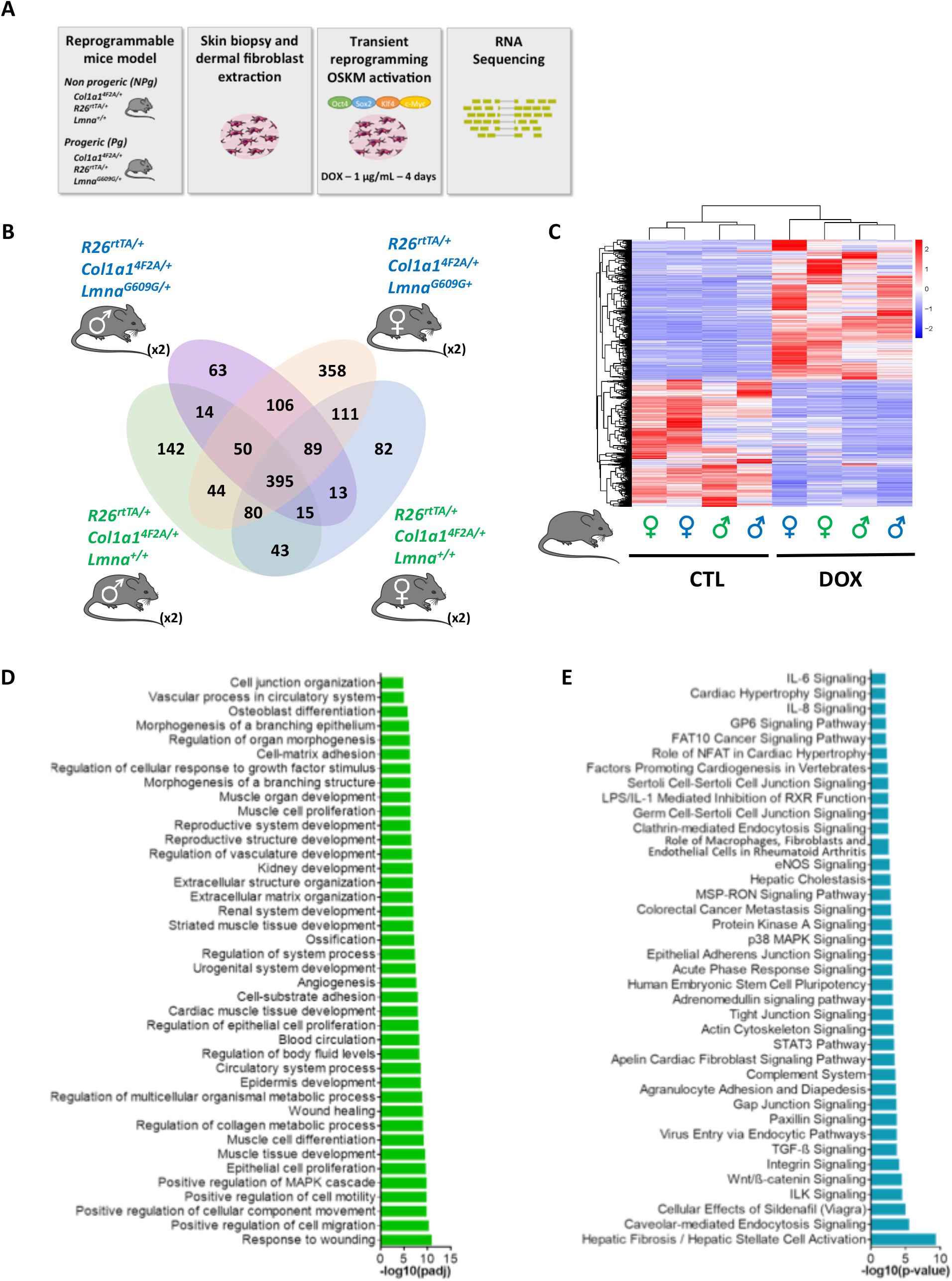
RNA sequencing reveals gene expression signature of *in vitro* OSKM induction independently of sexes and progeric genotype. **(A)** Schematic representation of the experimental procedures: dermal fibroblast were extracted from skin biopsies of 2 month-old R26^rtTA/+^;Col1A1^4F2A/+^;Lmna^G609G/+^ progeric and R26^rtTA/+^;Col1A1^4F2A/+^;Lmna^+/+^ non-progeric male and female mice. Expression of OSKM factors was induced *in vitro* by a daily exposure to doxycycline at 1µg/mL in culture medium for 4 days. Then, fibroblasts were harvested and RNA was extracted. **(B)** Venn Diagram presenting the number of differentially expressed genes in DOX samples versus CTL in progeric and non-progeric males and females. For each group, RNA from 2 mice were pooled for RNA sequencing. **(C)** Hierarchical clustering heatmap of the differentially expressed genes. Red color represents highly expressed genes and blue color represents low expression genes. Color descending from red to blue, indicates log10 (FPKM+1), from large to small. **(D)** Gene Ontology (GO) enrichment histogram displaying the top 10% significantly enriched (padj<0.05) biological processes common between all groups. **(E)** Top canonical pathways histogram revealed after analysis with Ingenuity software. The top 10% significantly affected pathways based on a restricted set of genes corresponding to the 395 differentially expressed genes common to all groups are highlighted.

Unexpectedly, GO and Ingenuity analysis identified that the major genes pathways targeted are consistent with formation or regeneration of organs and tissues, like cardiac and non-cardiac muscle tissue, kidney, bone but also with pathways related to diseases like fibrosis, dilated cardiopathy and osteoarthritis, as examples (Figure 3D, 3E). Although the experiment was performed in skin fibroblasts, gene pathways targeted are linked to general mechanisms involved in maintenance of tissues and organ integrity.

It incited us to investigate for the effect of the induction of OSKM on age related pathologies in our progeric mouse model.

### A single short reprogramming treatment ameliorates body composition and motor skills

To gain further insight of the impact of an early treatment on aging, we decided to study the potential effect of the single short reprogramming induction protocol on organismal metabolism and its consequences. Maintenance of lean mass and mobility is a pertinent indicator of health, both in human and mice (20). Firstly, we decided to follow the impact on body composition of our short early-in-life reprogramming protocol and secondly, we evaluated the impact on the age-related deteriorating mobility. Unexpectedly, while no significant effect on the total body weight evolution was revealed (Figure 4A), we observed a highly significant lower decrease of lean mass proportion in treated progeric animals, starting early after the treatment and maintained during aging (Figure 4B). Consistently, a highly significant decrease in the percentage of fat mass accumulation was observed suggesting that a global metabolic switch is triggered by our short reprogramming protocol, early in life, leading to the amelioration of body composition in treated animal (Figure 4B).

**Figure 4:**
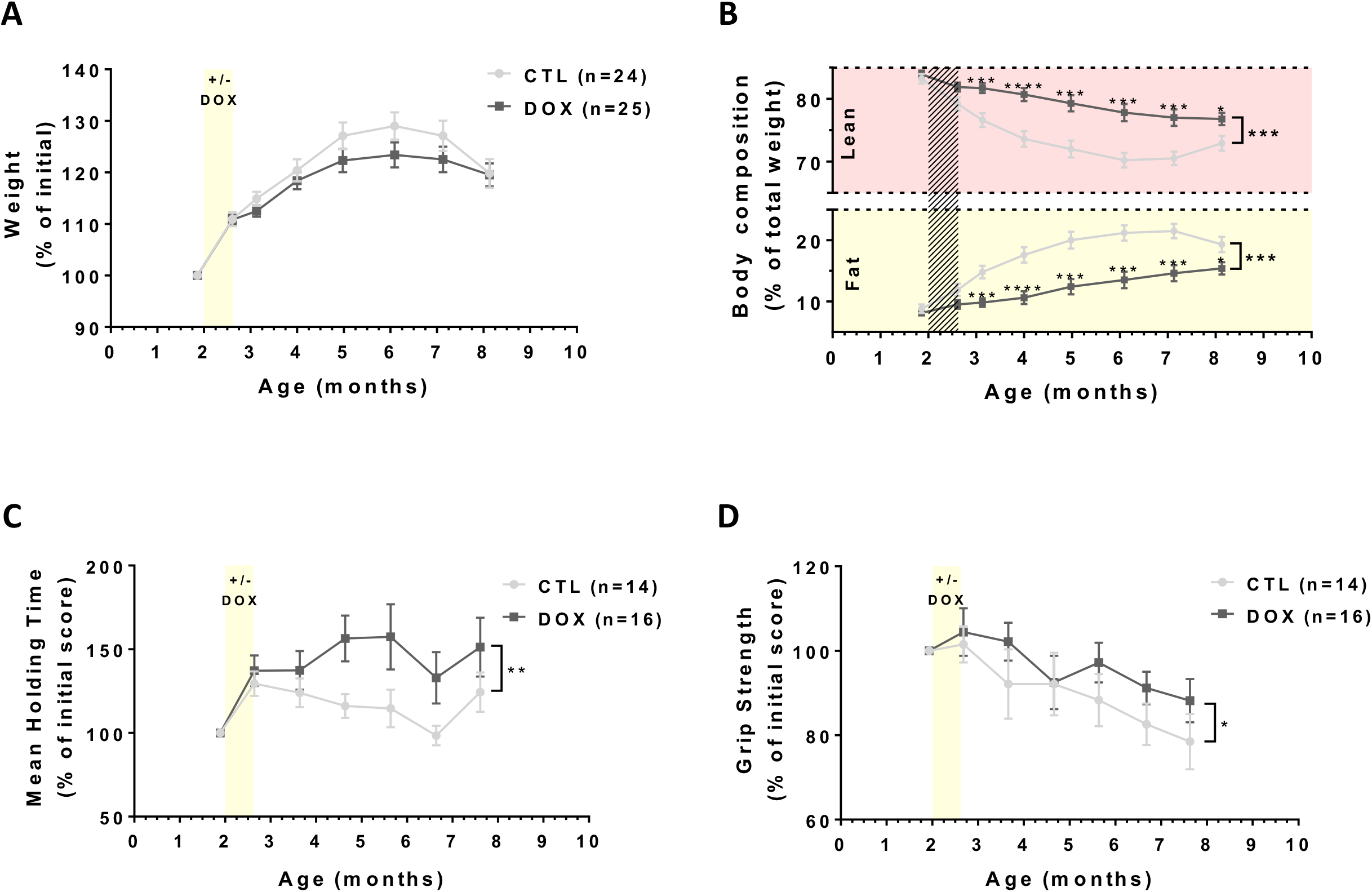
Early short transient reprogramming induces healthier body composition and improves lifelong muscular capacities in progeric mice. **(A)** Body weight curves of short-term treated R26^rtTA/+^;Col1A1^4F2A/+^;Lmna^G609G/+^ progeric mice compared with untreated controls. **(B)** Body composition through over life measured by EchoMRI-700. Results are expressed in percentage of total individual weight. **** p-value<0.0001; *** p-value<0.001; * p-value<0.05 according to multiple t-test for 1 vs 1 comparisons and paired t-test for whole curves. **(C)** Rotarod Assay. Maximal time to fall (strength endurance) compared to initial individual score. ** p-value<0.01 according to paired t-test. **(D)** Maximal Grip Strength compared to initial individual score. * p-value<0.05 according to paired t-test on whole curves.

As it generally favors healthy aging, we also evaluated the motor coordination of treated animals using the traditional increasing speed method on a rotarod (21) and we also tested muscle strength by grip tests. Both tests revealed improved motor skills in treated animal, initiated early after the treatment and maintained in aging (Figure 4C, 4D). These results demonstrate for the first time that a single short reprogramming induction early in life can initiate an immediate rejuvenation of body metabolism, whose positive consequences on motor skills are maintained during aging.

### A single short reprogramming treatment early in life ameliorates skin integrity in aging

As the largest organ of the body, skin, like many other tissues, gradually loses its self-renewal potential during aging and skin becomes thinner with age, losing its properties of resistance, plasticity and elasticity. This leads to a decrease of its capacity to optimally act as protective barrier and thus the organism becomes more vulnerable to external ionizing or microbial agents (22). Reduction in epidermal thickness is due to the reduction in the number of cells as well as to the degradation of the extracellular matrix as observed in histological sections from the elderly compared to the young (23). We thus analyzed skin structure by histochemistry. Strikingly we observed that our short reprogramming protocol with a dose of 0.5 mg/ml of doxycycline induced at two months of age led to a major protective effect on skin age-related thickness atrophy with a positive impact on all skin layers observed at 8 months of age (Figure 5A). There was a 40% average thickening of the epidermis and dermis, while the fat subcutaneous superficial layer and the panniculus carnosus smooth muscle layer increased by 120% (Figure 5B). Thus, this result demonstrates that the short reprogramming in the early life is able to delay skin deteriorations resulting in a maintained skin integrity at up to 8 months of age, suggesting that an engraved mechanism of this rejuvenating effect might be triggered by our short induction protocol.

**Figure 5:**
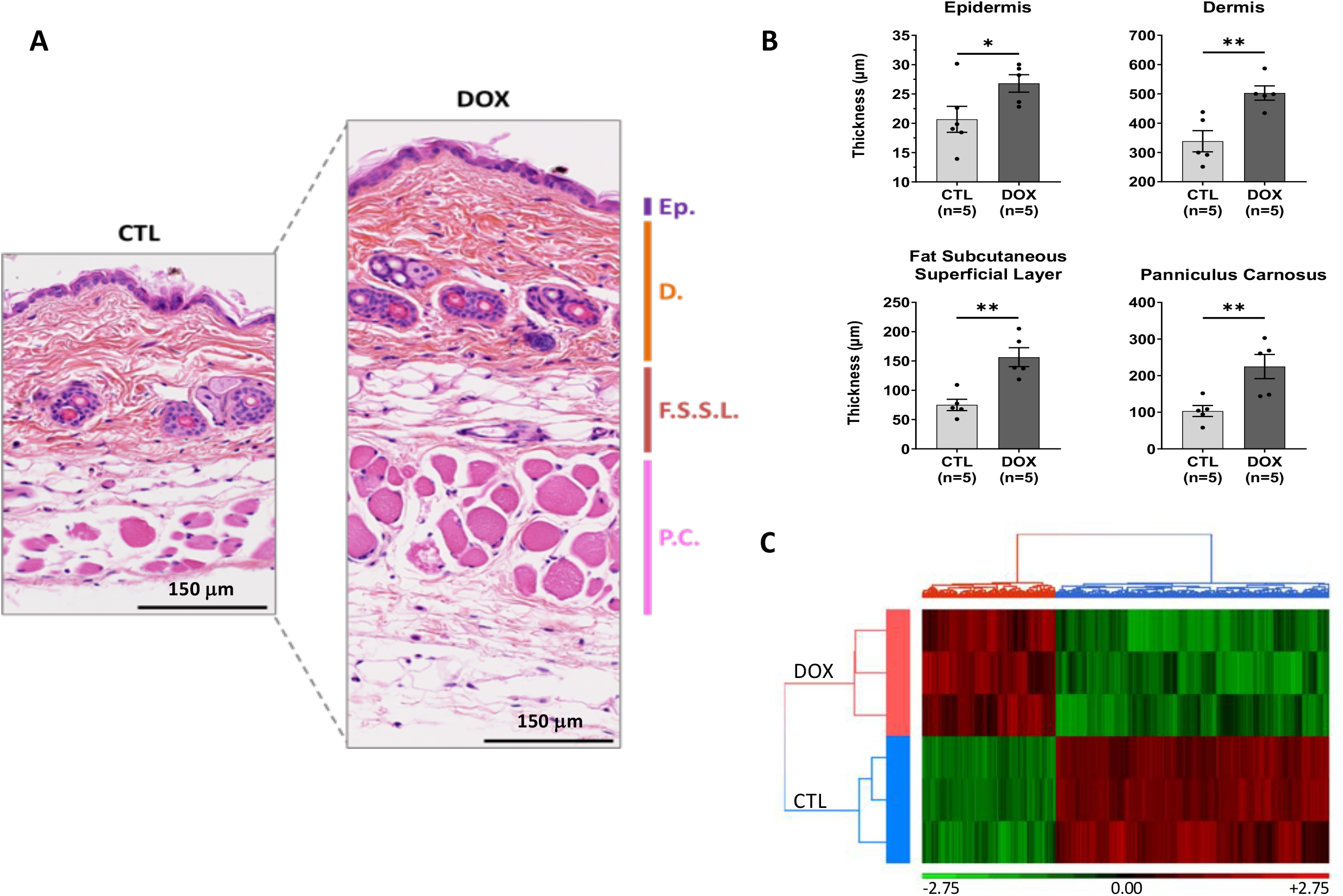
Early short transient OSKM induction induces amelioration of skin features in aging. **(A)** Histological analyzes of skin of 8 month-old R26^rtTA/+^;Col1A1^4F2A/+^;Lmna^G609G/+^ progeric mice. Skin layer thickness was measured after staining of skin section with Hematoxylin, Eosin and Saffron (HES). CTL represents untreated mice and DOX represents treated mice with 0.5mg/ml doxycycline over 2.5 weeks at the age of 2 months following our short-induction protocol. Ep.: Epidermis, D.: Dermis, F.S.S.L.: Fat Subcutaneous Superficial Layer, P.C.: Panniculus Carnosus. **(B)** Quantifications of layers thickness. All Measurements of areas and distances were performed on ImageJ software. **** p<0.0001; *** p<0.001; ** p<0.01; * p<0.05 according to unpaired t-test, two-tailed. **(C)** Hierarchical clustering of skin sample performed on significant (p<0.01) differentially methylated CpG loci between experimental mice control group (CTL, n=3) and doxycycline treated group (DOX, n=3). Loci Methylation levels of each CpG are represented by M value (log2 converted β values) which are shifted to mean of zero and scaled to standard deviation of one. Red and green colors represent respectively hypermethylated and hypomethylated CpG Loci. Doxycycline and control mice are respectively colored in red and blue.

### A single short reprogramming treatment triggers epigenetically related maintenance of skin integrity

Previous experiments using a reprogrammable mouse model demonstrated that a cyclic induction of OSKM two days a week, over the entire lifetime of a homozygous accelerated aging mouse model, increased longevity, eventually through a chronical epigenetic remodeling of unstable histone marks (12). We thought that our short induction protocol might engraved more permanent marks of the “rejuvenation effect” that could explain the maintained integrity of skin observed in aging. Aging evokes dynamic changes in DNA methylation (DNAm) at specific CG dinucleotides (CpG) (24), as a biomarker for the aging process, often referred to as “epigenetic clock” (3). DNA methylation is described to reflect aspects of biological age since the deviation of predicted and chronological age correlates with all-cause mortality in human (4). Consequently, modifications in DNA methylation seemed pertinent to investigate for such a “memorized effect” initiated by our short induction protocol early in life.

Various “epigenetic clock” for mice were initially described on Whole Genome Bisulfite Sequencing from multi-tissue specimens and were clearly demonstrated to be affected by genetic, dietary, or pharmacological interventions (5-7). Recently, Illumina Infinium Mouse Methylation arrays covering 280 000 CpG sites, were released allowing us to investigate for differentially methylated regions across the mice genome. In skin samples from 8 months-old mice, normalized methylation level was calculated for each CpG locus and a one-way-ANOVA statistical test reveals that 2641 differentially methylated CpG loci allowing to cluster control (CTL, n=3) and doxycycline treated (DOX, n=3) mice groups (Figure 5C). Although it does not exist an available “epigenetic clock” designed from these Mouse Methylation arrays for our progeria model, our results indicate that our short reprogramming protocol induced early in life might engraved a DNA methylation related epigenetic memory promoting an amelioration skin function in aging. Interestingly, GO analysis with the 70 genes associated to this 2641 differentially methylated sites, revealed that 26 genes and miRNAs are related to aging (Supplementary Table 1). Interestingly, some of them are directly or indirectly involved in chromatin remodeling (25), DNA damage response pathways (26), stress resistance (27) and longevity (28) and some miRNAs like mir125b-1 are preferentially expressed by skin stem cells regulating self-renewal and differentiation (29). Altogether, these results suggest strongly that our short induction protocol might improve the biological age of the skin through an epigenetically related mechanism.

### Amelioration of tissue structure and function at the onset of age-related pathologies

Because several tissues are frequently altered in the elderly, leading to the initiation and progression of age-related diseases, we tested multiple tissues and organs for reduced degeneration (25). To this end, tissues and organs of mice treated at two months of age, according to our single short reprogramming protocol with a dose of 0.5 mg/ml of doxycycline were sampled at the age of 8 months and analyzed.

#### A single short reprogramming treatment prevent age related osteoarticular diseases

The first pathology associated with aging that we studied on our model was osteoarthritis (Figure 6). In humans, it is the most common joint disease. At the knee level, it is characterized by an alteration of all the structures of the joint and in particular by a degradation of the cartilage which can even go as far as a degradation of the subchondral bone. The etiology of the pathology is a complex combination of hereditary and environmental determinants (30). Aging is one of the main factors, with an increasing prevalence of osteoarthritis over the years. We thus wanted to assess whether there was an improvement in cartilage and bone tissue in the knees of our treated mice. We studied the cartilage of the lateral and median plates of the left knee by confocal laser scanning microscopy (CLSM) and the subchondral bone of these same plates was analyzed by X-ray microtomography (µ-CT). The CSLM measurements revealed that the mice induced at the age of 2 months had a significantly higher volume of cartilage and a reduced cartilage surface degradation 6 months later, when compared to untreated control mice (Figure 6A). This trend was confirmed in the subchondral bone by a decrease in the degradation of the surface exemplified by a smoother appearance of the bone on the 3D reconstruction (Supplementary Figure 4A). Thus, these results suggest that a single short reprogramming induction early in life induces a long-lasting protective effect against knee cartilage degradation.

**Figure 6:**
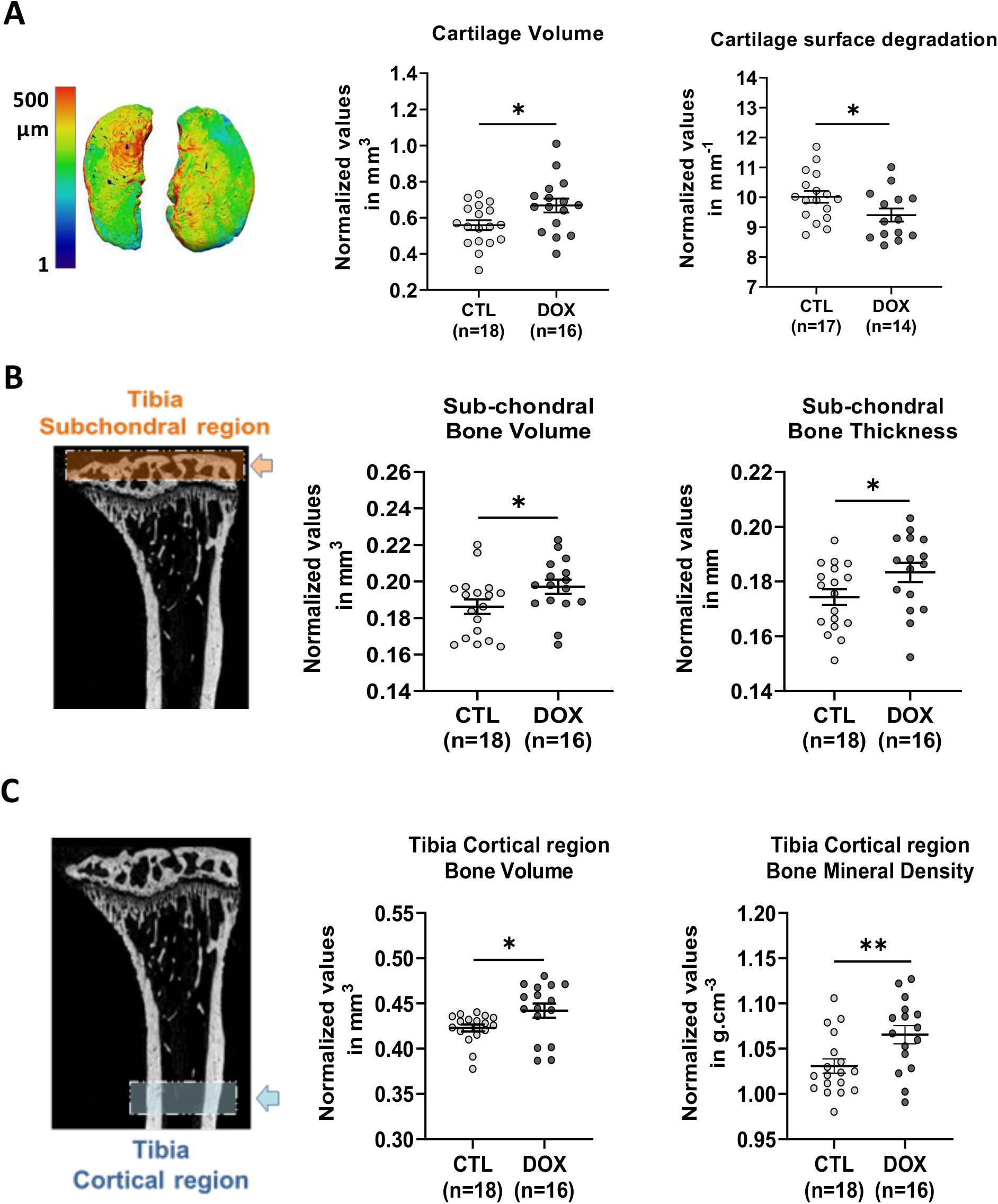
A single short OSKM induction early in life prevents osteoarthritis and osteoporosis in aged mice. **(A)** Histomorphometric analysis of 3D images of knee joint cartilage by confocal laser scanning microscopy (CLSM). Cartilage volume, and surface degradation were measured in the lateral and medial plateau. * p<0.05 according to unpaired t-test, two-tailed. **(B)** X-ray micro-computed tomography (μ-CT). Histomorphometric analysis of left tibia sub-chondral bone, in the knee joint (orange box and arrow). Sub-chondral bone volume and thickness were measured in the lateral and medial plateau. **(C)** µ-CT histomorphometric analysis of tibia cortical region (blue box and arrow). Cortical bone volume and mineral density were measured on both tibias. Bone and cartilage tissues were analyzed on 8 month-old R26^rtTA/+^;Col1a1^4F2A/+^;Lmna^G609G/+^ progeric mice. CTL represents untreated mice and DOX represents treated mice with 0.5mg/ml doxycycline during 2.5 weeks at the age of 2 months following short-induction protocol. For µCT analysis, ** p<0.01; * p<0.05 according to unpaired t-test, one-tailed, with Welch’s correction.

Another bone pathology commonly encountered during aging is osteoporosis. This skeletal pathology decreases bone mass and deteriorates its internal structure making it more fragile, and greatly increases the risk of fracture (31). The causes of osteoporosis are manifold and still mostly unknown. However consequences of subchondral bone structure alterations associated to osteoporosis might be an early event in osteoarthritis pathology (32). As with osteoarthritis, the prevalence of osteoporosis increases with age and although it affects both sexes there is a two to three times higher frequency in women due to menopause and the decrease in estrogen. To assess the level of osteoporosis in our animals, we firstly analyzed the subchondral bone subjected to alterations in microstructure in aging. We observed, a larger volume as well as a greater thickness of the bone (Figure 6B). Then, we analyzed the cortical region of two tibia by µ-CT. We were able to observe a higher bone volume and mineral density in the induced mice without any difference in bone thickness nor in its surface degradation, suggesting a lower porosity in the treated mice compared to the controls (Figure 6C, Supplementary Figure 4B).

Collectively these results demonstrated that a single short induction of cell reprogramming factors early in life positively regulates aging features of bone by protecting from osteoarthritis and osteoporosis in later life.

#### A single short reprogramming treatment prevent age related tissue structure deteriorations and fibrosis

With a view to deepen our understanding of the effects of the transient reprogramming, enabled by our short treatment, we extended our analysis to a wide panel of organs to highlight macroscopic changes in their structure and integrity. During the aging process tissue integrity gradually deteriorates under the action of numerous cellular and molecular events such as the exhaustion of stem cells, present in the tissues and of their capacities to differentiate for tissue renewal, as well as the accumulation of senescent cells and tissue fibrosis. Fibrosis, a pathological process characterized by aberrant inflammatory injury and excessive fibrous connective tissue production (33), is accepted as the primary cause for age-related deterioration of various human organs, including lung (34), kidneys (35), liver (36) and heart (37). Although little is known about the molecular mechanisms involved, accumulation of senescent cells and the Senescence Associated Secretory Phenotype (SASP), is thought to be a common cause of multiple age related pathologies (38).

Idiopathic pulmonary fibrosis, is a severe pulmonary impairment, most often resulting in death, 2 to 5 years after diagnosis, from respiratory failure. Most patients referred with idiopathic pulmonary fibrosis are over 70 years of age. Indeed, age, male sex and smoking are important risk factors for the development of this condition (39, 40). Idiopathic fibrosis results in excessive accumulation of extracellular matrix and remodeling of lung architecture, in particular with the filling of the alveolar space with connective tissue. To explore the impact of our short reprogramming on lung alterations and fibrosis, we used the validated Sirius Red or Masson’s Trichrome staining. Quantification of lung fibrosis, in mice treated by our short early-in-life reprogramming protocol, showed significant decrease of covered areas at 8 months of age (Figure 7A).

**Figure 7:**
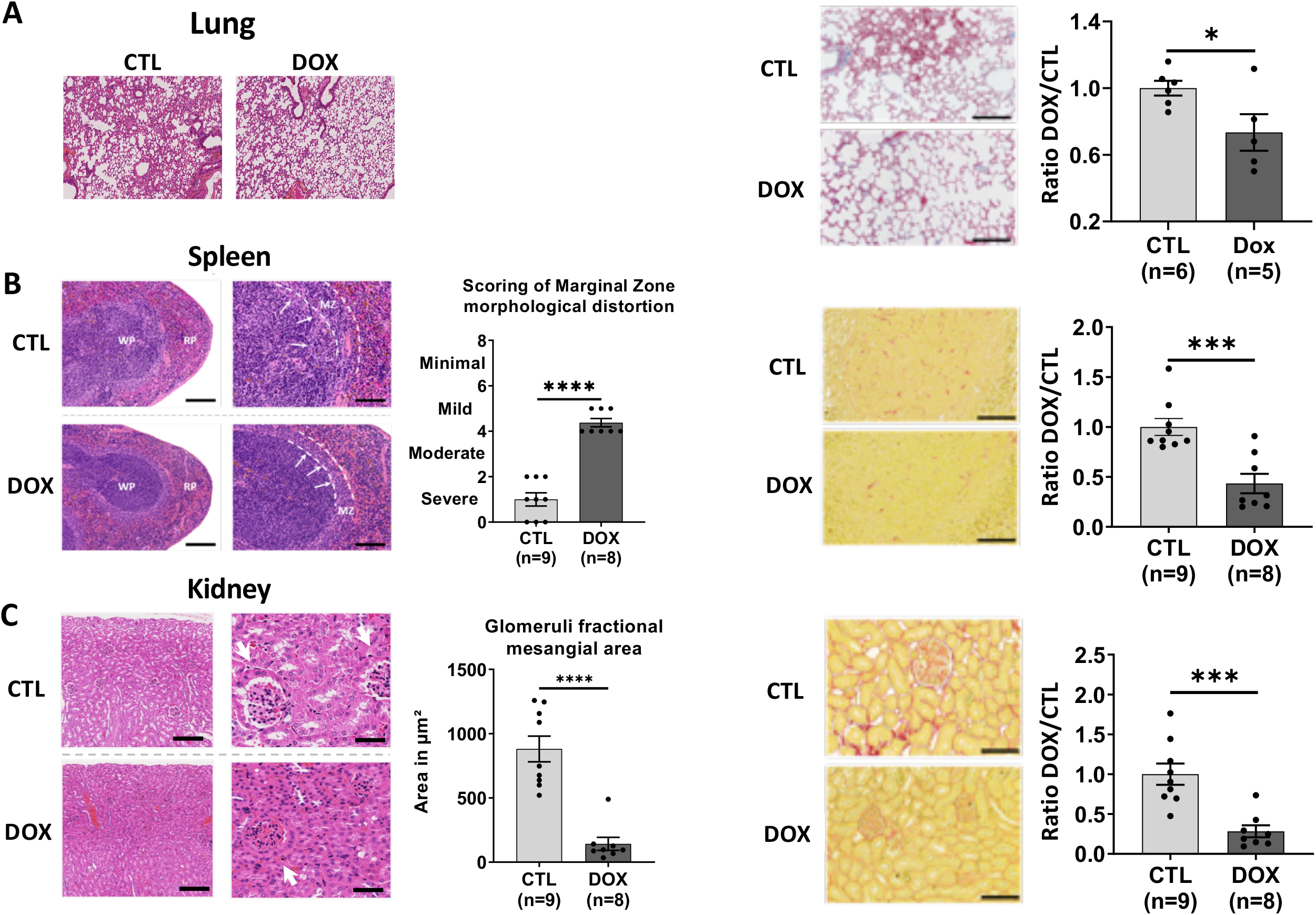
Tissue structure and age-related tissue fibrosis is improved in aging by a single OSKM induction early in life. Histological analyzes of lung, spleen and kidneys sections were performed after staining with Hematoxylin, Eosin and Saffron (HES) and fibrosis after Red Sirius (RS) and Masson’s Trichrome (MT) staining. Both staining was used to detect collagen fibers and stained areas were measured after color deconvolution, as described in methods section. **(A)** Morphologic comparison of lung structure and fibrosis in treated and untreated mice. Scoring of fibrosis level as described in methods. **(B)** Morphologic comparison of the Marginal Zone (MZ) architecture of spleen from treated and untreated mice. The MZ/White pulp interface distortion is depicted by the inner line, and the percent radius involvement (MZ protruding into the white pulp area) is depicted by arrows. Scoring was determined as described (58) and of the spleen white pulp fibrosis was scored as described in the methods section. **(C)** Measurement of kidney’s fractional mesangial area, representing the space surrounding glomeruli (depicted by arrows) and inter-tubular fibrosis. All tissues were analyzed on 8 month-old R26^rtTA/+^;Col1a1^4F2A/+^;Lmna^G609G/+^ mice. CTL represents untreated mice and DOX represents treated mice with 0.5 mg/ml doxycycline for 2.5 weeks at the age of 2 months following short-induction protocol. All Measurements of areas and distances were performed on ImageJ software. **** p<0.0001; *** p<0.001; ** p<0.01; * p<0.05 according to unpaired t-test, two-tailed.

Spleen structural loss of integrity is commonly observed in the elderly. Spleen, as well as lymph nodes, is an important secondary lymphoid organ involved in immune response to pathogens and prevention of senescent cells accumulation during aging (41). Thus, structural disorganization of the spleen during aging may compromise the immune system. Consequently, this immune-senescence leads to decreased response to vaccination and increased susceptibility to infectious diseases in aged patients. Histological analysis of the spleen revealed significant changes between our two groups of animals. Indeed, after scoring the architecture of the marginal zone, we determined that the treated animals obtained an average score ranging from 4 to 6, characterizing a mild alteration against an average score ranging from 0 to 2 characterizing a severe alteration of the marginal zone in the control animals (p<0.0001, Figure 7B). We have thus highlighted a greater maintenance of the marginal zone structure, at the interface between the white and red pulp, in animals that have been transiently reprogrammed.

Kidneys are also affected during aging where a loss of integrity and increased fibrosis alters their function, and may result in an increased susceptibility to drug toxicity and a potentially harmful electrolyte imbalance. The histological features of kidneys from the elderly include decreased cortical mass, glomerulosclerosis, interstitial fibrosis, tubular atrophy, and arteriosclerosis. Our study of the renal tissue first focused on the space surrounding the glomeruli, also named the fractional mesangial area and which is known to increase during aging (42). In the group of mice treated with doxycycline we measured a strong significant decrease of the fractional mesangial area of 85% on average (p<0.0001, Figure 7C) with an associated decreased fibrosis (Figure 7C).

Thus, in addition to the amelioration of organ structure, we concluded that a global decrease of fibrosis at the organismal level was observed and particularly significant in lungs, spleen and kidneys with an additional trend in liver and heart (Supplementary Figure 5). This might be beneficial to healthspan.

With this latter result, our data clearly demonstrated that structure and integrity of tissues, in older individuals, is positively impacted by a single short reprogramming, through the expression of OSKM, early in life.

Since we showed previously that this induction protocol fixed an epigenetic memory, through DNA methylation, it incited us to further analyze this process on the different organs studied.

### Amelioration of tissue structure and function at the onset of age-related pathologies by an epigenetically related mechanism

Like previously in skin, we analyzed DNA methylation on the various organs studied, spleen, kidney, lung, heart and liver. We identified the best differential methylation signatures specific for each organ in clustering control (CTL, n=3) and doxycycline treated (DOX, n=3) mice groups by using a one-way-ANOVA statistical test (p-value >0.01) (Figure 8A). The best CpG signature identified in each case seems to be very specific of each organ, as they differ from one organ to another in terms of CpG and associated genes loci (Figure 8B, Supplementary Table 2).

**Figure 8:**
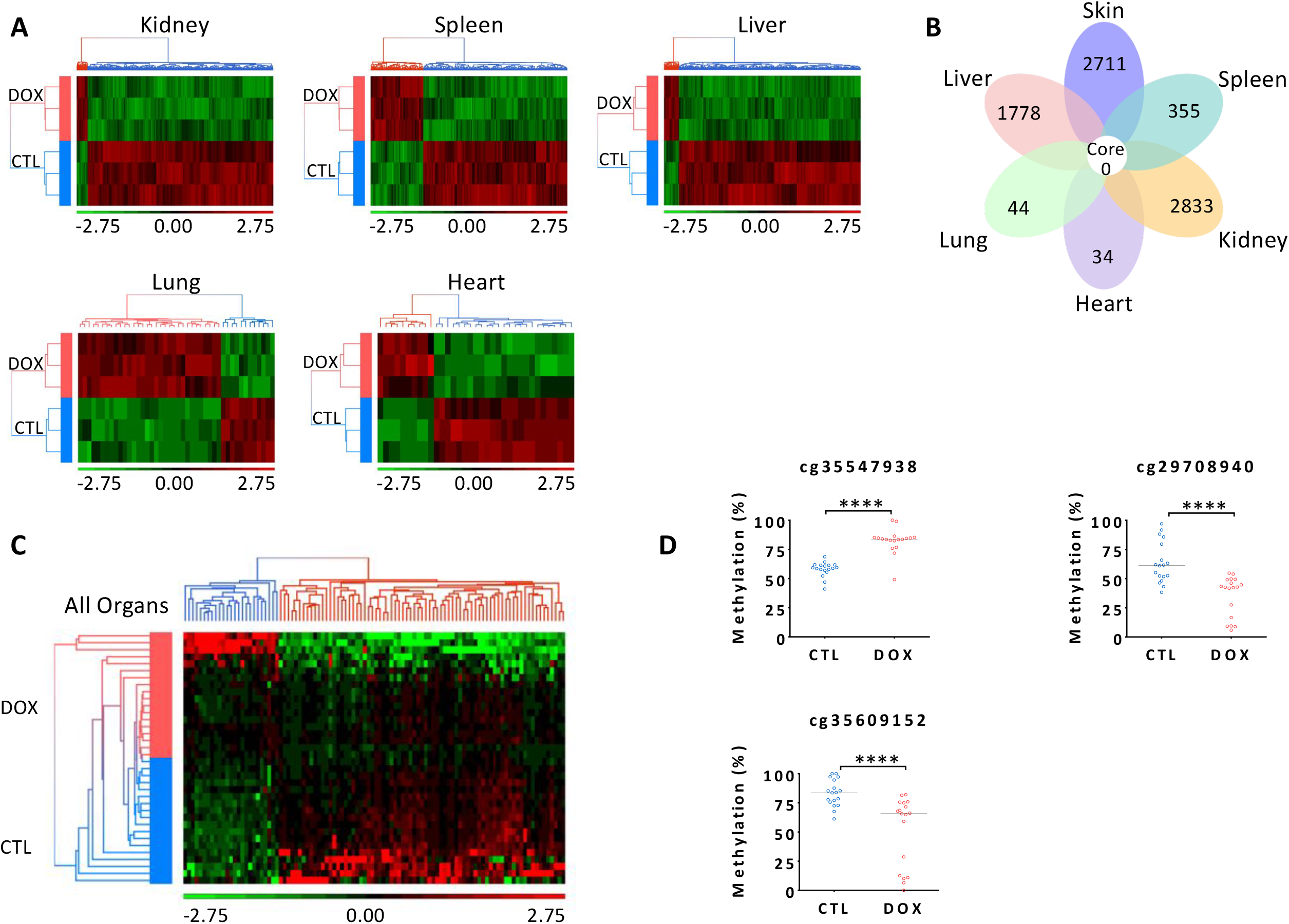
Differential methylation analysis on mice reveals a sustainable signature set by a single short OSKM induction early in life. **(A)** Hierarchical clustering tissue by tissue performed on significant (p<0,01) differentially methylated CpG loci between experimental mice control group (CTL, n=3) and doxycycline treated group (DOX, n=3). Loci methylation levels of each CpG are represented by M value (log2 converted β values) which are shifted to mean of zero and scaled to standard deviation of one. The Red and green color represent respectively hypermethylated and hypomethylated CpG Loci. Doxycycline and control mice are respectively colored in red and blue. **(B)** Flower diagram on each tissue signatures. List of differentially methylated CpG is compared organ by organ (DOX vs CTL with p<0.01). Each petal represents a tissue with the number of differentially methylated CpG. The core shows the number of shared CpG between the 6 tissues signatures. **(C)** Hierarchical clustering performed on 100 most significant differentially methylated CpG loci between experimental mice control group (CTL, n=18) and doxycycline treated group (DOX, n=18) for all tissues. Two successive statistical tests were used to select the 100 loci list. One-way ANOVA (p<0.05) followed by Mann-Whitney statistical test was applied. The hundred most significant Mann-Whitney value were selected (p<0.0005). Loci methylation levels of each CpG are represented by M value (log2 converted β values) which are shifted to mean of zero and scaled to standard deviation of one. The Red and green color represent respectively hypermethylated and hypomethylated CpG Loci. Doxycycline and control mice are respectively colored in red and blue. **(D)** Scatter plot performed on CpG loci corresponding to the 3 highest methylation differences between control (CTL, n=18) and doxycycline (DOX, n=18) mice group among “all organs’ signature”. Loci methylation levels of each CpG (β values percentage) are compared between experimental mice control group (CTL, n=18) and doxycycline treated group (DOX, n=18). All individuals’ tissues methylation levels are plotted. Doxycycline and control mice are respectively colored in red and blue and median β values of each group are represented by a black line. Significance of differences between control and doxycycline treated mice are indicated (Mann-Whitney test: **** p<0.0001).

To further obtain a signature that could reflect the decreased physiological age of our treated mice, we included all the organs studied in the analysis and then we selected the 100 loci with the highest difference in methylation between control and doxycycline treated groups (Figure 8C, Supplementary Table 3). This highly significant signature reveals that our short induction protocol delivered at 2 months is able to memorize DNA methylated regulations correlated with a positive impact on functions of various tissues at 8 months of age.

In an attempt to develop a pan tissue “epigenetic clock” on our progeria model, we selected loci, based on their β values, corresponding to the 3 highest methylation level differences between control (CTL, n=18) and doxycycline (DOX, n=18) mice groups (Figure 8D; Supplementary Table 4). We propose that this specific signature extracted for the 280 000 loci might allow in the future to calculate the biological age of our treated mice to evaluate the extent and propagation of the rejuvenated physiology triggered by our single short OSKM reprogramming protocol.

## Discussion

In this paper, we developed a specific model of heterozygous progeric mice as an intermediate model, between wild-type and severely affected homozygotes, to study rejuvenating interventions. The heterozygotes live about 35 weeks compared to 20 weeks for homozygotes and 2 years for wild type. The heterozygotes are thus very sensitive to improvements while not being so afflicted as to be difficult to interpret. Ocampo and collaborators developed a specific induction protocol showing that by inducing a short and chronic induction two days a week of the Yamanaka reprogramming factors OSKM through the entire life of an accelerated-aging mouse model, they extended the lifespan of mice and improved some age-related hallmarks (12). However, the use of a homozygous model of severe accelerated aging where the mice live for only 20-or-so weeks in standard breeding settings questions the physiological relevance of the observed phenomenon. We thus decided to refine and simplify the Ocampo protocol using heterozygotes progeric mice and we explored the impact of a short transient reprogramming on aging and age-related pathologies in longer-living animals. Although various local reprogramming induction protocols were described on different mice models to study the immediate impact on tissue fitness, they led to direct or indirect beneficial or deleterious effect (43-50). Here, we discovered that a single step of cellular reprogramming at the level of the organism by a two and half weeks of treatment on two month-young heterozygotes impacts both lifespan and healthspan, protecting tissues and organs that deteriorate during aging.

The long-delayed benefits of the early short OSKM induction, lead us to conclude that the triggered immediate cell rejuvenating effect we observed, initiates protective molecular mechanisms in tissues, as suggested by our RNAseq analysis, in which selecting common genes differentially expressed in progeric and non-progeric cells by a short induction of OSKM, identified similar gene pathways involved in tissue and organ integrity mechanisms.

Ocampo et al. proposed that the effect observed is epigenetically related, but also demonstrated that a chronic induction of OSKM two days a week, over the entire lifetime on its homozygous accelerated aging mouse model, is necessary to maintain the epigenetic remodeling followed by unstable histone marks (12). To overcome this potential epigenetic instability, we developed a protocol with an increased period of induction of a single two and a half weeks instead of two days to engrave more permanently the marks of a rejuvenated cell physiology In consequence, we observe a “memorized effect” at the onset of increased tissue deterioration in aging, which is associated to a comprehensive DNA methylation signature shared by all the organs studied. Although, we cannot define precisely the mechanism involved, the fact that this single induction period early in life promotes beneficial effects on the global physiology in aging suggests that the DNA methylation changes triggered by OSKM induction might be involved in the initiation and propagation of this rejuvenated cell physiology. Although further experiments will be needed, this indisputable ‘distal’ effect, ultimately turn out to increased lifespan and healthspan through an epigenetic reprogramming.

Previously, we added a further two factors to the OSKM cocktail and observed reversal of senescence *in vitro* as an additional effect (51). As a combination of 6 Factors OSKMNL can rejuvenate old cells *in vitro* by either total or partial reprogramming (11, 51), it will be also very interesting to test if the effect observed *in vivo* by a treatment early in life with OSKM might be amplified by the OSKMNL combination. The six factors cocktail might equally be efficient when applied on aged mice in reversing the aging phenotype.

Population aging, and more importantly healthy aging, is a major public health issue with important economic and social impact. Consequently, identifying pharmaceutical targets or therapeutic strategies would help to increase lifespan and more specifically health span (52, 53). Nowadays, clearing senescent cells accumulating in aging is the most promising strategy envisaged to promote healthy aging (54). It has been demonstrated on progeric and non-progeric mice in seminal works on a genetically modified model triggering apoptosis in p16 expressing cells (55, 56). This proof of concept allowed the development of several senolytic molecules selectively triggering apoptosis in senescent cells in clinical trials to treat age related pathologies. Also, although we clearly do not envision a purely genetic clinical approach to induce a short reprogramming with OSKM, the results we have obtained are a proof of concept that by unique short reprogramming even early in life is sufficient to promote healthy aging.

This therefore open the door to potential clinical applications, using molecules mimicking this early memorized rejuvenating treatment to delay senescence and to erase aging and age-related diseases, either alone or in combination with senolytic treatments (57).

## Methods

### Mice model and housing

Mice allowing *in vivo* transient reprogramming were originally developed by Rudolf Jaenisch (Whitehead Institute, Massachusetts Institute of Technology, USA) and purchased from Jackson Laboratories (STOCK Gt(ROSA)26Sor^tm1(rtTA+M2)Jae^ Col1a1^tm3(tetO-Pou5f1,-Sox2,-Kfl4,-Myc)Jae^/J (JAX: 011004). These mice are bred in homozygous form for the two transgenes (two copies for the rtTA transactivator (R26^rtTA/rtTA^) and two copies for the 4 reprogramming factors (OSKM) cassette (Col1A1^4F2A/4F2A^). The murine line exhibiting the accelerated aging phenotype (Hutchinson-Gilford progeria syndrome) was originally developed by Carlos Lopez-Otin from University of Oviedo, Spain (Lmna^tm1.1Otin^ (MGI: 5295747))(14). This line carries the G609G mutation on the LMNA gene, leading to the activation of a cryptic splicing site and the accumulation of the prelamin A truncated form, also called progerin. This line is bred in heterozygous form for the progeria mutation (Lmna^G609G/+^). Experimental groups were generated by the crossing of these two lines and are represented by the following genotypes: progeric R26^rtTA/+^;Col1A1^4F2A/+^;Lmna^G609G/+^ and non-progeric R26^rtTA/+^;Col1A1^4F2A/+^;Lmna^+/+^.

Animal care and use for this study were performed in accordance with the recommendations of the European community (2010/63/UE). The Project was validated by the Ethical committee of the French Ministry of Research through the agreement APAFIS #21760. All mice were produced at Plateau Central d’Elevage et d’Archivage du CNRS de Montpellier (PCEA, Agreement n°A34-172-45) and transferred to the analysis platform one week before starting experiments for habituation. The procedures and protocols concerning body composition and functional analysis were performed on the Metamus platform, at the DMEM unit of the INRAE-UM in Montpellier, France (Veterinary Services National Agreement n° E34-172-10, 04 March 2019). All the others procedures and protocols were performed at INM (Agreement n° C34-172-36). Mice were housed in groups in filter-top cages with free access to standard diet (A04, SAFE diets, Augy, France) and tap water. They were maintained in a temperature-controlled room (24°C ±1°C) in a standard 12:12 light-dark cycle (lights at 7:30 am). All cages were enriched with nesting materials (cellulose squares, SAFE). Animal behavior was checked daily for welfare and health status and mice weight was monitored every two weeks. For mice subjected to body composition analysis and functional assays, cages were also enriched with a hanging red polycarbonate tunnel. This tube is equivalent to the animal holder used for EchoMRI measures and allows mice habituation to a restrained environment.

### Longevity studies

To induce reprogramming in our animals, we implemented either lifelong or short induction protocols. For both, doxycycline was administered in drinking water in opaque bottles. All protocols started at the age of two months and were carried out on animals of genotype R26^rtTA/+^;Col1a1^4F2A/+^;Lmna^G609G/+^ or R26^rtTA/+^;Col1a1^4F2A/+^;Lmna^+/+^. Two lifelong protocols were used and lasted until the animals died. The first one that we have developed consists of a lower induction at 0.2 mg/ml of doxycycline, maintained continuously every day of life. In addition to this, two short induction protocols were developed. Both consists in inducing animals for only 2.5 weeks at the age of two months, at 0.2 mg/ml of doxycycline for the first and 0.5 mg/ml for the second. The last one was also developed on non-progeric animals of genotype R26^rtTA/+^;Col1a1^4F2A/+^;Lmna^+/+^.

### Body composition analysis

Mice whole-body composition (fat and lean masses) was measured every month throughout the study by quantitative magnetic resonance with a whole-body composition analyzer (EchoMRI™-700 Echo Medical Systems, Houston, TX, USA) according to the manufacturer’s instructions. Mice were individually weighted before each measurement.

### Functional Assays

To measure the motor coordination of mice, we used a rotarod machine with automatic timers and falling sensors (47650 Rota-Rod, Ugo Basile®). We set up the machine in ramp mode from 5 rpm to 40 rpm over 300 seconds. Before acquiring the initial data, 2 months old mice were accustomed and trained once on the rotarod machine. Then they performed a test once a month until time of death. All mice were placed on the 3 cm diameter fluted cylinder, the run started at 5 rpm for a few seconds until all mice were facing the running direction and then the rotating speed increased. We recorded the latency to falling of each animal. The animals were placed back in their cages directly after falling from the cylinder. Each animal ran 3 times for a maximal time of 10 minutes followed by 20 minutes of resting between each run.

To assess the muscular strength of mouse front legs, we performed a vertical grip strength test (Bio-GS3, BioSeb). Before acquiring the initial data, 2 months-old mice were accustomed and trained once on the machine. Then they performed a test once a month until time of death. Mice were held by the tail and placed above the gripping bar until they grasped the bar. Then the animal was pulled up until grip was released. We recorded the maximal grip force developed. Each animal was submitted to the test 3 times, with a few minutes rest, in its cage between each test. The test was always performed by the same experimenter.

## Acknowledgements

We thank Biocampus Montpellier for technical support and expertise as well as all the staff involved for animal housing on RAM (Réseau des Animaleries de Montpellier) facilities, for metabolism experiments on RAM-Metamus platform at the DMEM laboratory, for tissue processing on RHEM (Réseau d’Histologie Expérimentale de Montpellier) facility, on MRI (Montpellier Ressources Imagerie) platform for imaging, on the National platform ECELLFrance for µ-CT and CLSM analysis and the PPM (Pôle Protéome de Montpellier) platform for metabolomic analysis. We thank Metamontp platform supported by the European Regional Development Funds (ERDF) for EchoMRI experiments. We thank Dr Julian Venables for the edition of the manuscript.

## Authors contributions

Conceptualization and Supervision: J.M.L., O.M. Funding acquisition and project administration: J.M.L. Data analysis: Q.A., E.L.B., P.B., O.M., J.M.L. Writing original draft: Q.A., E.L.B., P.B., O.M., J.M.L. Animal experiments and sample analysis: Q.A., E.L.B., C.L., O.M. Histology: Q.A. In vitro experiment: Q.A., E.L.B. RNAseq: Q.A., E.L.B. Methylome: P.B. Animal model: C.L. Bone and cartilage: Q.A., K.T., D.N., C.J. Proteomic: E.L.B., J.V., C.H. Animal metabolism profiling: E.L.B., C.B.G., L.P., F.C. Scientific and technical support: N.B., M.G., F.E.

## Data availability

The datasets generated and analyzed during the current study are available from the corresponding authors on reasonable request.

## Supplemental informations

**Supplementary Figure 1:**
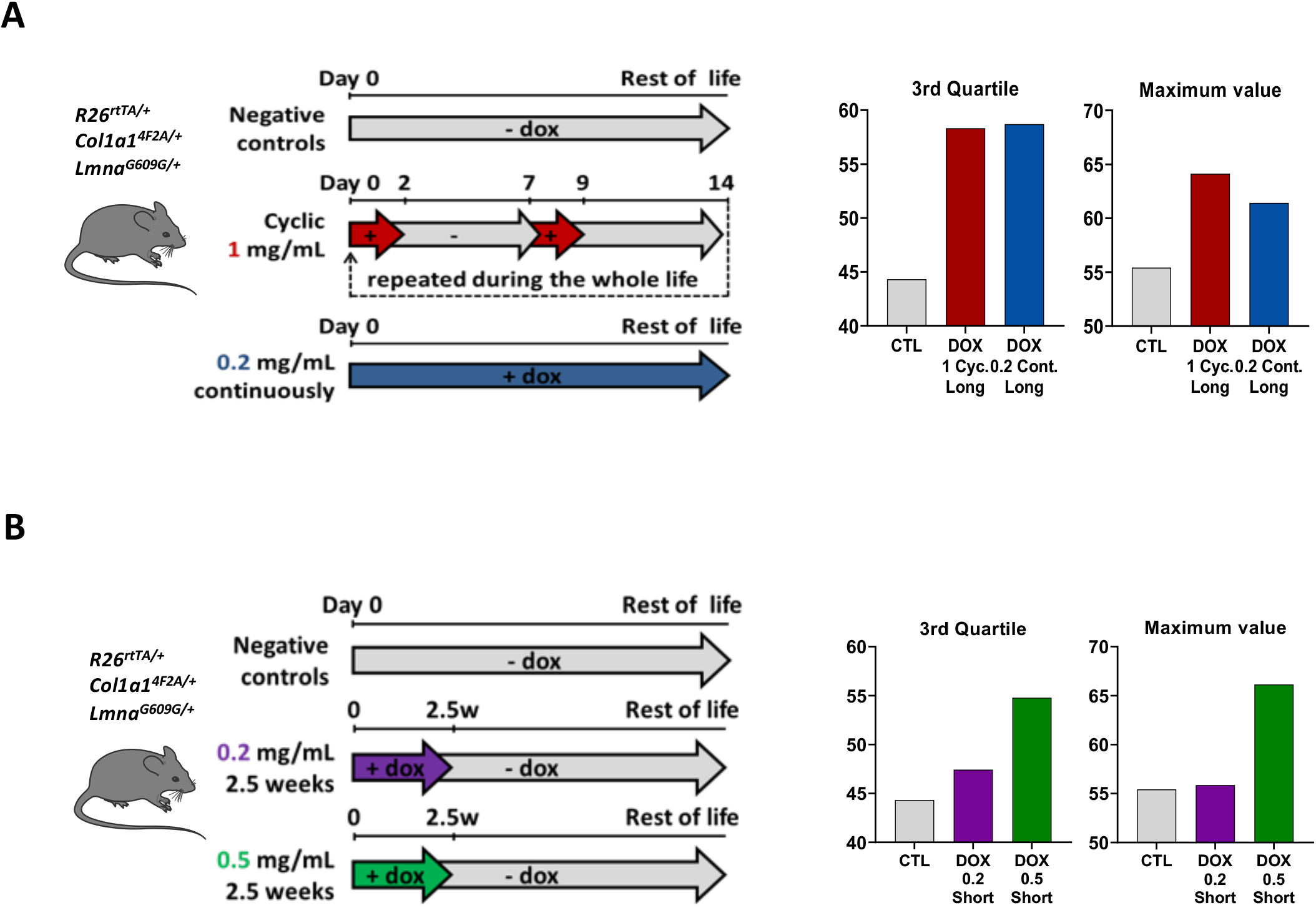
Longevity parameters of continuous and single short OSKM induction protocols. **(A)** Scheme of lifelong OSKM doxycycline induction protocols for R26^rtTA/+^;Col1a1^4F2A/+^;Lmna^G609G/+^ progeric mice. Median survival for the 3^rd^ quartile and maximum lifespan value are presented. **(B)** Scheme of the single short OSKM doxycycline induction protocols for R26^rtTA/+^;Col1a1^4F2A/+^;Lmna^G609G/+^ progeric mice. **** p<0.0001; ** p<0.01 according to unpaired t-test, two tailed. Median survival for the 3^rd^ quartile and maximum lifespan values are presented.

**Supplementary Figure 2:**
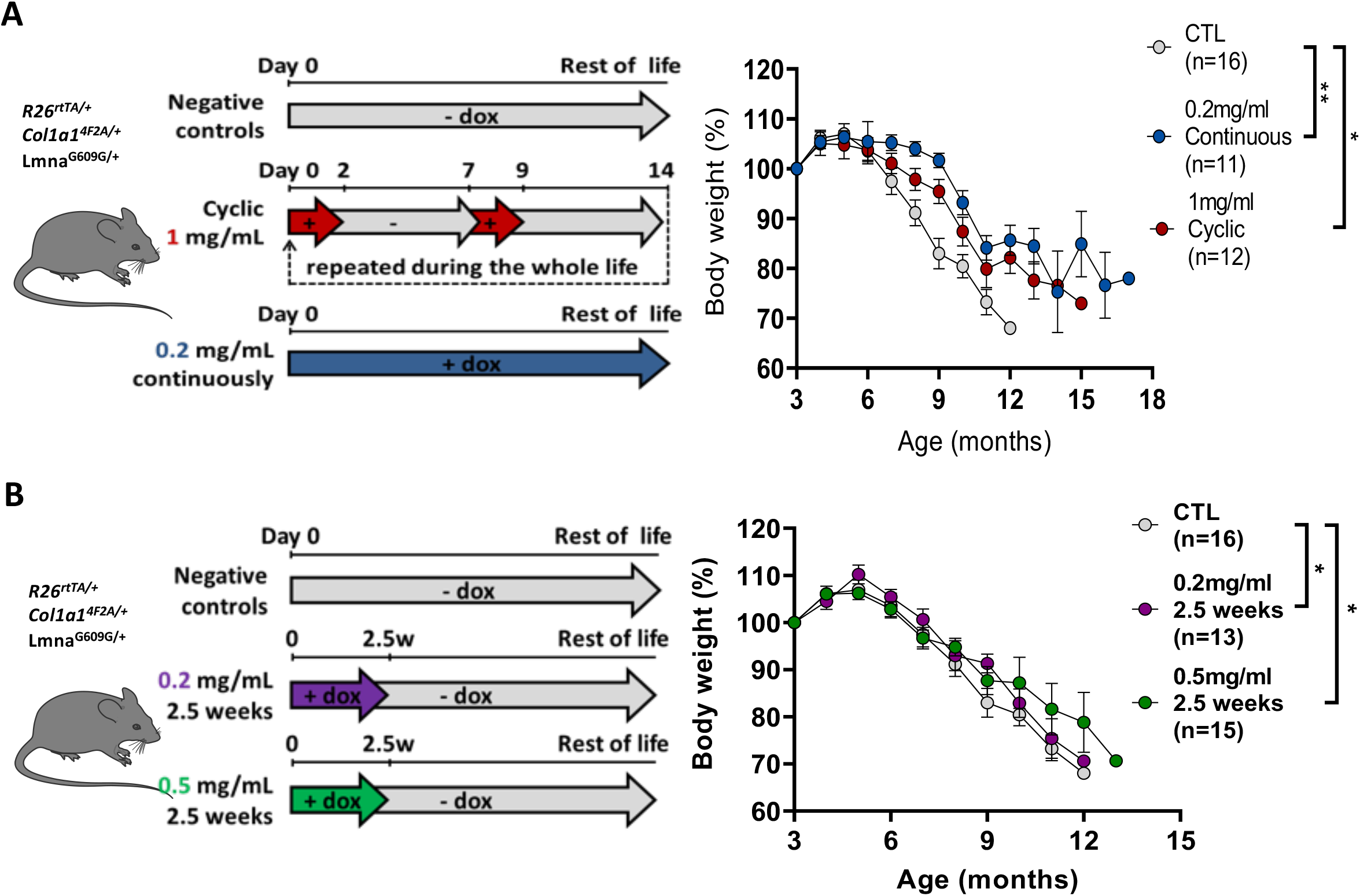
Effect of continuous and single short OSKM induction protocols on weight loss. **(A)** Scheme of lifelong doxycycline induction protocols for R26^rtTA/+^;Col1a1^4F2A/+^;Lmna^G609G/+^ progeric mice. Body weight curves of treated mice compared with untreated controls. ** p<0.01; * p<0.05 according to paired t-test, two tailed. **(B)** Scheme of single short OSKM doxycycline induction protocols for R26^rtTA/+^;Col1a1^4F2A/+^;Lmna^G609G/+^ progeric mice. Body weight curves of treated mice compared with untreated controls. * p<0.05 according to paired t-test, two tailed.

**Supplementary Figure 3:**
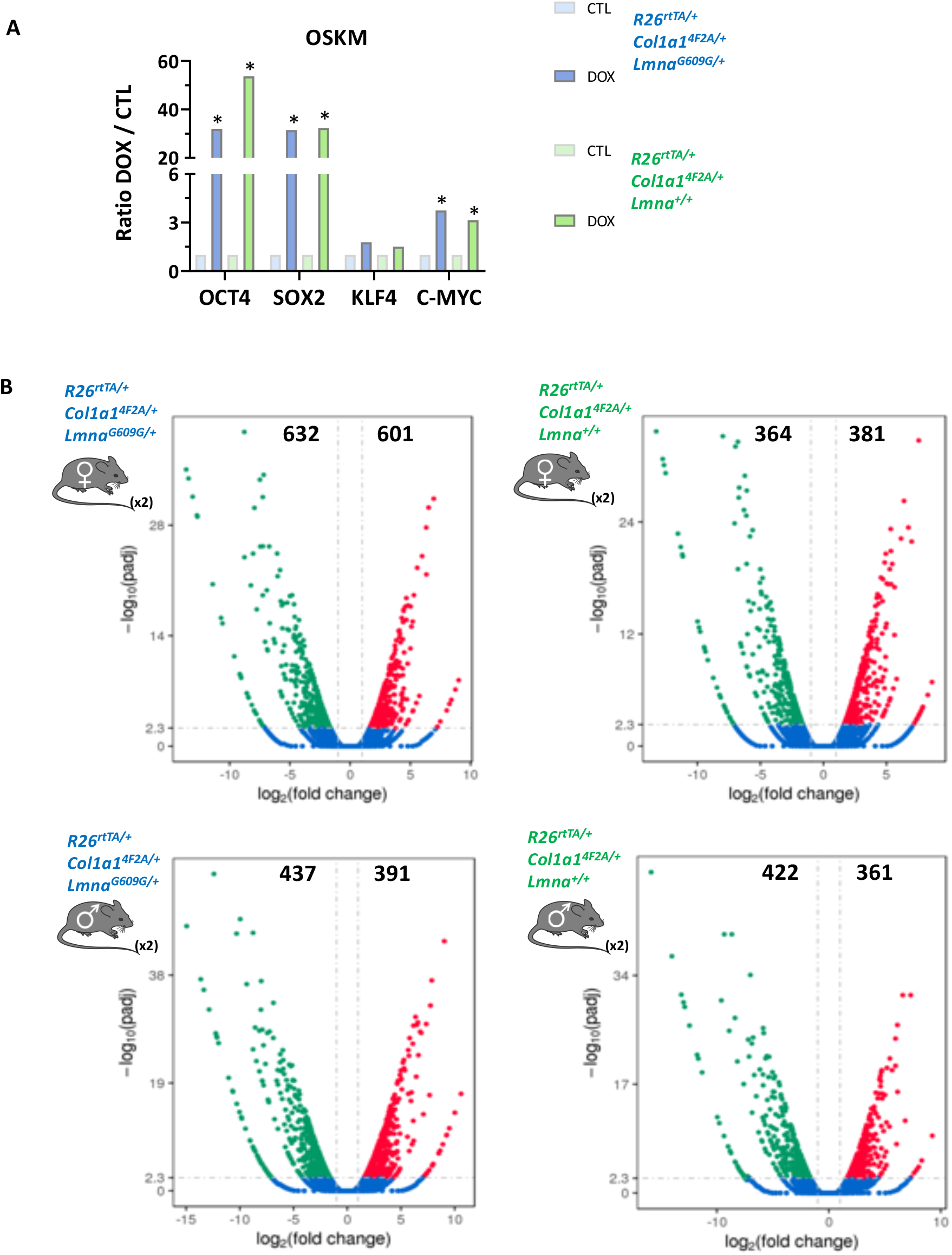
Differential gene expression analysis in skin fibroblasts induced by OSKM. **(A)** Expression levels of OSKM factors in *in vitro* induced skin fibroblasts of progeric and non-progeric, female and male mice. **(B)** Volcano plot displayed the differential fibroblast-induced gene expression from progeric and non-progeric males and females. Number of genes up and down regulated are mentioned. Expression of OSKM factors was induced *in vitro* by a daily exposure to doxycycline at 1µg/mL and volcano plot of differential fibroblast-induced gene expression from progeric and non-progeric males and females. Number of genes up and down regulated are mentioned.

**Supplementary Figure 4:**
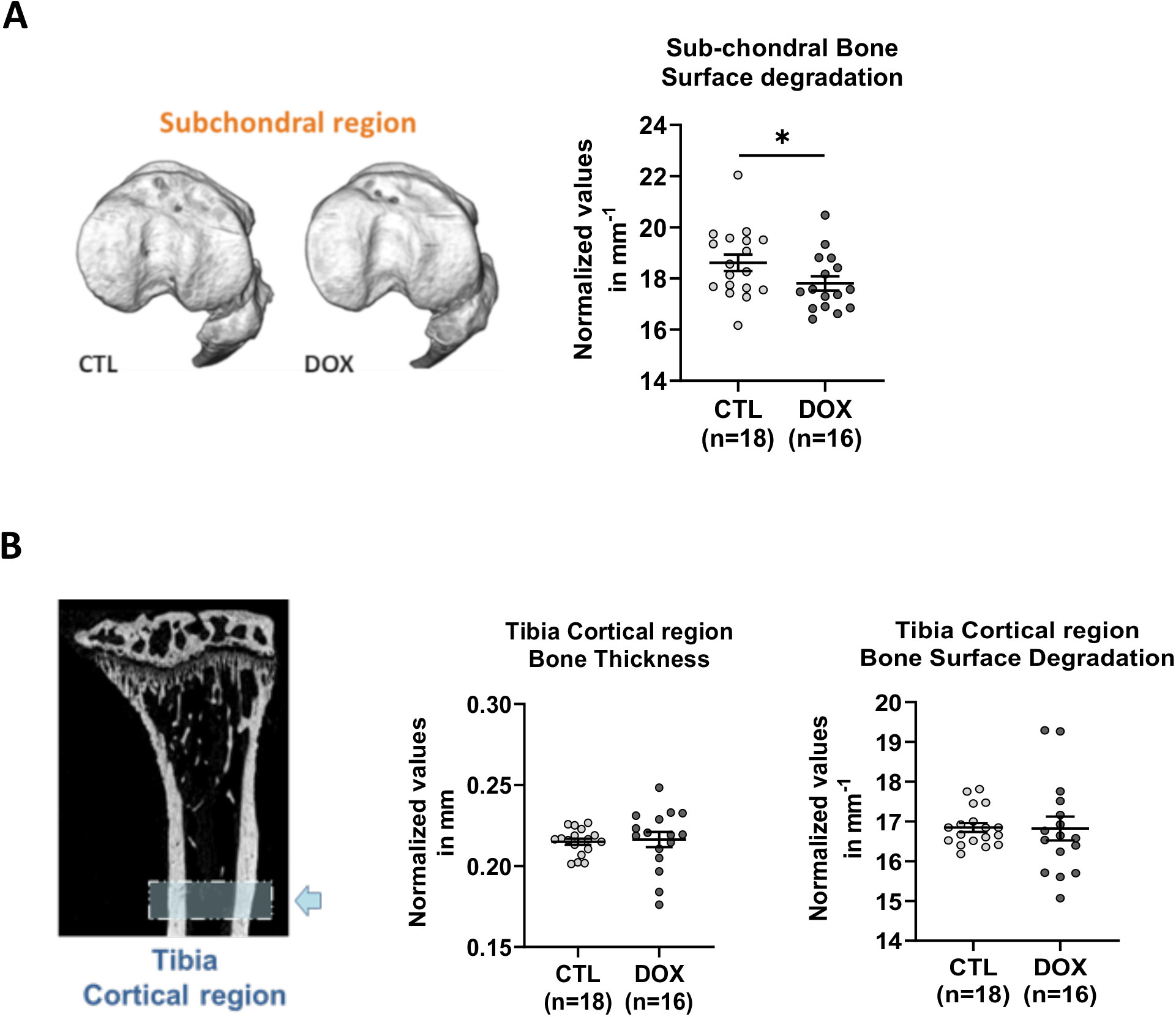
A single short OSKM induction, early in life, prevents osteoarthritis and osteoporosis in age mice. **(A)** Representative 3D reconstruction of sub-chondral bone is represented on the upper right corner of the panel. Rough appearance is representative of bone degradation. Sub-chondral bone surface degradation was measured in the lateral and medial plateau. **(B)** µ-CT histomorphometric analysis of tibias cortical region (blue box and arrow). Cortical bone thickness and surface degradation were measured on both tibias. Bone tissues were analyzed on 8 months old *R26*^*rtTA/+*^; *Col1a1*^*4F2A/+*^;Lmna^G609G/+^ progeric mice. CTL represents untreated mice and DOX represents treated mice with 0.5 mg/mL doxycycline during 2.5 weeks at the age of 2 months following short-induction protocol. * p<0.05 according to unpaired t-test, one-tailed, with Welch’s correction.

**Supplementary Figure 5:**
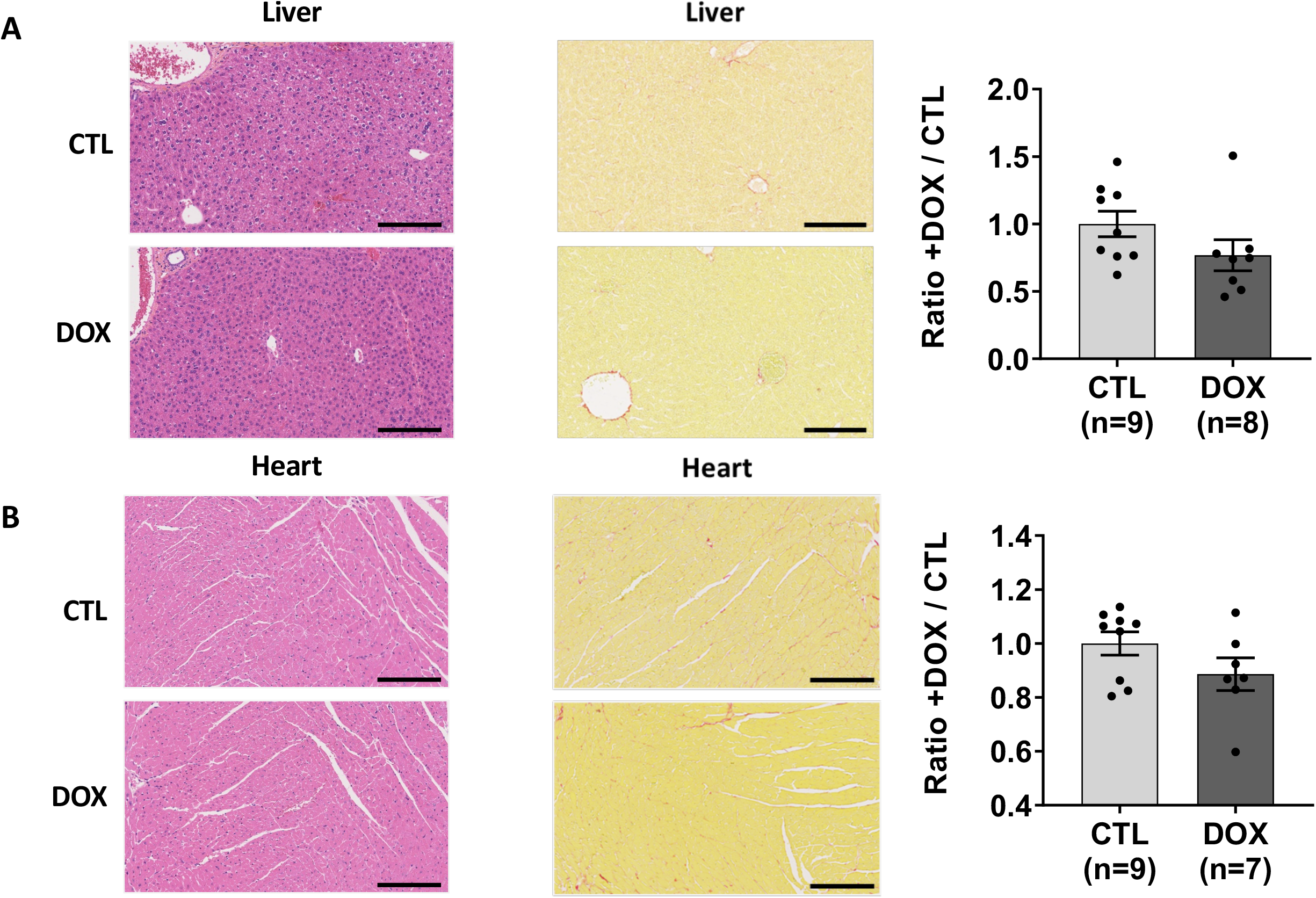
Tissue structure and age-related tissue fibrosis is improved in aging by a single OSKM induction early in life. Histological analyzes of Liver **(A)** and heart **(B)** sections were performed after staining with Hematoxylin, Eosin and Saffron (HES) and fibrosis after Red Sirius (RS) and Masson’s Trichrome (MT) staining. Both staining was used to detect collagen fibers and stained areas were measured after color deconvolution as described in methods section. **(A)** Morphologic illustration of liver tissue structure and fibrosis analysis in treated and untreated mice. **(B)** Morphologic illustration of heart left ventricle tissue structure and fibrosis analysis in treated and untreated mice. All tissues were analyzed on 8 months old R26^rtTA/+^;Col1a1^4F2A/+^;Lmna^G609G/+^ progeric mice. CTL represents untreated mice and DOX represents treated mice with 0.5 mg/ml doxycycline during 2.5 weeks at the age of 2 months following short-induction protocol. All Measurements of areas and distances were performed on ImageJ software. **** p<0.0001; *** p<0.001; ** p<0.01; * p<0.05 according to unpaired t-test, two-tailed.

**Supplementary Table 1: List of genes associated to the differentially methylated loci and belonging to aging Go term in skin**.

The table summarizes the genes associated to the differentially methylated loci in skin and among them all genes belonging to Aging Go term that are found by the global Gene Ontology analysis performed on significant (p<0,01) differentially methylated CpG loci between experimental mice control group (CTL, n=3) and doxycycline treated group (DOX, n=3). “Ensembl Transcripts release 100” was chosen as annotation database. Promoter region was configured as 5000 base pairs upstream and 3000 base pairs downstream from the transcription start site (TSS).

**Supplementary Table 2: List of genes associated to the differentially methylated loci and belonging to aging Go term by tissue**.

The table summarizes the genes associated to the differentially methylated loci tissue by tissue and among them all genes belonging to Aging Go term that are found by the global Gene Ontology analysis performed on significant (p<0,01) differentially methylated CpG loci between experimental mice control group (CTL, n=3) and doxycycline treated group (DOX, n=3). “Ensembl Transcripts release 100” was chosen as annotation database. Promoter region was configured as 5000 base pairs upstream and 3000 base pairs downstream from the transcription start site (TSS). All unique genes are listed in tissue’s corresponding excel files.

**Supplementary Table 3: List of the 100 selected CpG loci for all organs**.

The table summarizes the hundred most significant loci selected for their highest methylation differences between control (CTL, n=18) and doxycyclin (DOX, n=18) mice group among “all organs’ signature” (see Figure C).

**Supplementary Table 4: List of genes associated to the 3 best differentially methylated loci**.

The table summarizes the genes associated to CpG loci corresponding to the 3 highest methylation differences between control (CTL, n=18) and doxycycline (DOX, n=18) mice group among “all organs’ signature” (see Figure 8D). “Ensembl Transcripts release 100” was chosen as annotation database. Promoter region was configured as 5000 base pairs upstream and 3000 base pairs downstream from the transcription start site (TSS).

## Supplementary Methods

### Isolation of dermal fibroblasts

Dermal fibroblast were extracted by enzymatic dissociation from skin biopsies of 2 months-old progeric R26^rtTA/+^;Col1A1^4F2A/+^;Lmna^G609G/+^ and non-progeric R26^rtTA/+^;Col1A1^4F2A/+^;Lmna^+/+^ mice. For each animal, the shaved back skin was removed after euthanasia, cleaned with Betadine (povidone-iodine) and rinsed with cold 1X PBS before being removing. The skin was decontaminated with three successive antibiotics solution baths (MEM Hepes, Fungizone, Gentamycin) then cut into square fragments (0.5cm × 0.5xm) in a clean cell culture dish. The pieces were covered with a Dispase (Roche) solution (HBSS, Dispase II 2.5U/ml, Soybean 0.1%) and incubated overnight at 4°C.

Using forceps, dermis and epidermis from each piece were separated and placed in two different 50 ml tubes. Afterwards, 9 ml of trypsin/EDTA were added to each tube and this was placed at 37°C for 20 min, with shaking every 5 min (gentle for the epidermis and strong for the dermis). The tubes were then passed through a 100 μm sieve, pieces were recovered for a second trypsination and then discarded. The trypsin was then inactivated by adding 10% FBS (calcium free for the epidermis) and the cells were mechanically dissociated 20 times with a 5 ml pipette. The suspensions were then passed through a 70 μm sieve and 10 ml of DMEM were used to rinse the sieves. The cells were counted and centrifuged for 5 min at 300 g. The fibroblasts were then resuspended and cultured in DMEM glutamax medium, 10% FBS, 1% PenStrep, 1% NEAA (Gibco).

### Gene expression profile by RNA sequencing

Expression of OSKM factors was induced *in vitro* by a daily exposure to doxycycline at 1µg/mL in culture medium for 4 days. Then, fibroblasts were harvested and RNA extraction was performed with a RNeasy Mini Kit (Cat No. 74104, QIAGEN) according to manufacturer recommendations. RNA was further processed by Novogene Corporation for RNA sequencing. Briefly, after qualification of RNA and library construction, libraries were fed into an Illumina sequencer NovaSeq 6000. Raw data were then mapped to the mouse reference genome and differential gene expression analysis and enrichment analysis (GO, KEGG) were performed.

### Western blot analysis

Lysis of tissue and cell samples was performed on ice in RIPA buffer (Fisher Scientific) supplemented with HALT protease and phosphatase inhibitor cocktail (Fisher Scientific). Samples were then centrifuged at 14,000 g for 15 min at 4°C to remove the cellular debris. Then, protein concentration in lysates was quantified using the BCA protein assay kit (Fisher Scientific).

For western blot experiments, NuPAGE gels and related devices and reagents (Fisher Scientific) were used. The protein lysates were denatured for 10 min at 70°C in LDS buffer with reducing agent and then loaded into 4-12% or 12% Bis-Tris Novex polyacrylamide gels. The migration was carried out in MOPS buffer for 45 min at constant 200v. The transfer was carried out in transfer buffer containing 20% EtOH and 1 ml of antioxidant, for 1 hour at constant voltage (30v).

For each experiment, the transfer membranes were rinsed for 5 min with 1X TBS before the blocking step for 1 hour with 10% non-fat dry milk in 1X TBS. Immunoblotting was performed in a 1X TBST buffer with 1% BSA. Primary antibodies were incubated overnight at 4°C. The membranes were washed 3 times with 1X TBST and incubation with the secondary antibody was performed for 1 hour at room temperature. The membranes were washed twice with 1X TBST and then 1 time with 1X TBS before revelation. Visualization was carried out with the SuperSignal West Pico Chemiluminescence Substrate kit (Fisher Scientific) and revealed using a ChemicDoc reader (Bio-Rad) and the Image Lab 4.1 software, which was used for quantification.

### Immunofluorescence

All cells were seeded on 22 × 22mm glass coverslips gelatinized with a 0.1 % gelatin solution (Millipore) for a minimum of 30min. The cells were washed twice with 1X PBS and fixed with 2% formaldehyde in 1X PBS for 10 min at RT. Then, the fixative solution was replaced by 1X PBST, avoiding any drying. For permeabilization, cells were incubated in 0.5% Triton-100 in 1X PBS for 10min at RT. The non-specific epitopes were blocked with a 1X PBS solution with 2% BSA and 0.5% fish skin gelatin for 30 min. Primary antibodies were incubated for 1 h in a humid chamber at room temperature. Unbound antibodies were removed by 3 thorough washes in 1X PBST. The secondary antibodies were incubated for 1 h in a humid chamber at room temperature, in the dark. Then, the DNA was stained with DAPI (2 μg/mL) for 10 min and the excess was washed 3 times with PBST and twice with MilliQ water. The samples were mounted on a SuperFrost glass slide with Mowiol and sealed with nail polish. All the antibodies used were diluted in blocking solution buffer. Sample acquisitions were realized using Zeiss Axiovert 200M equipped with HXP120V light source under AxioVision Rel.4.8 software.

### Antibodies

Peroxidase-conjugated AffiniPure Goat Anti-Mouse IgG (Jackson ImmunoResearch 115-035-146; WB: 1/50000). Peroxidase-conjugated AffiniPure Goat Anti-Rabbit IgG (Jackson ImmunoResearch 111-035-144; WB: 1/50000). Mouse mAb anti-GAPDH (Ambion AM4300 clone 6C5; WB: 1/10000; Lot 095P055535A). Rabbit pAb anti-H3 (Abcam ab 1791; WB: 1/5000; Lot 172452). Rabbit pAb anti-LC3B (Sigma-Aldrich L7543; WB: 1/1000, IF 1/500; Lot 102M4778V). Rabbit mAb anti-Oct-4A (Cell Signaling Technology #2840 clone C30A3; WB: 1/1000; Lot 15 Ref 04/2016). Rabbit mAb anti-γH2AX Ser139 (Cell Signaling Technology #9718 clone 20E3; WB: 1/1000; Lot 12 Ref 08/2015). Rabbit mAb anti-Fox03a (Cell Signaling Technology #2497 clone 75D8; WB: 1/1000; Lot 6 Ref 04/2016). Goat pAb anti-rabbit IgG Alexa Fluor 488 (Molecular Probes, Ref: A11034 Lot: 1531670).

### Mitochondrial DNA quantification

After 4 days of OSKM induction, cells were harvesting and DNA extraction were performed using DNAzol Reagent according to manufacturer recommendations (Cat No. 10503027, Invitrogen™). Quantified and qualified DNA samples were then used as template for qPCR (Light Cycler 480, Roche) in order to quantify the mtDNA copy number by determining the mtDNA/nDNA ratio, according to previously described method (59).

### Metabolomics

To extract metabolites secreted within the cells, cells were harvested and pellets were immediately suspended in Methanol/CAN 1/1 solution. After centrifugation for 10 minutes at 13,000 rpm, supernatants were collected and stocked at -80°C until analysis. 10µL of a standard solution (pyruvic acid 13C1 and succinic acid 3H4 at 2.5 mM, Sigma Aldrich, Saint Louis, USA) was added to 50µl of supernatant. Sample was dried under nitrogen (50°C) and resuspend with 100µL of formic acid 0.1 % (Biosolve, Dieuze, France). 10µL of the sample was injected on an UPLC system (Acquity, Waters Corporation, Milford, USA) coupled to a triple quadrupole mass spectrometer (Xevo TQD, Waters Corporation) was used for targeted metabolomic assay in MRM mode. RPLC separation was performed with an ACQUITY UPLC HSS T3 (2.1×100 mm, 1.8 mm, Waters Corporation) column. The mobile phase A was water (Biosolve, Dieuze, France) with 0.1% formic acid (v/v) and mobile phase B was methanol (Biosolve, Dieuze, France) with 0.1% formic acid (v/v) with an optimized gradient-elution program: 0-0.2 min, 99% A; 0.2-1.0 min, 98%-10% A; 1.0-2.0 min at 10 % A, 2.0-2.3 min 10-99 %A; 2.3-3.5 min at 99 % A at a flow rate of 400µL/min. The column temperature was maintained at 45°C. The key parameters for MS ion source were set as follows: Negative ESI mode, cone tension 2.5kV, desolvatation temperature 500°C, cone temperature 150°C, desolvatation flow rate 800L/h, and the cone flow rate 50L/h.

The optimized MRM parameters were transitions 86.77>43.02, 88.77>43.02 114.90>70.90, 116.90>72.90, 132.90>70.90, 190.96>86.90 to pyruvic, lactic, fumaric, succinic, malic and citric acid respectively and m/z 87.77>43.02, 120.80>77.07 to pyruvic acid 13C1 and succinic acid 3H4 respectively.

### Tissue sampling and preparation

In these experiments, animals of the short-term OSKM induction group (0.5 mg/ml of doxycycline at 2 months for 2.5 weeks and untreated controls) were sacrificed at 8 months following an ethical and authorized procedure, 6 months after the end of the induction. Nine controls animals and eight treated animals with both sexes were used in these studies.

Organs were removed and dissected according to RITA standardization procedures(60). The samples intended for tissue structure analysis were fixed in 4% PFA, for 24 h at room temperature, then washed three times in 1X PBS and stored in 70% EtOH at 4°C before staining.

At the same time, tissue samples were also snap-frozen in liquid nitrogen for protein, RNA and genomic DNA analysis. The samples were placed in 1.5 ml tubes with stainless steel beads of different sizes (1 × 3mm, 2 × 2mm and 4 × 1mm in diameter) and ground in an adequate buffer using the Qiagen MixerMill MM300 tissue lyser. More details are available in dedicated sections.

For cartilage and bone analysis, whole hind legs were fixed in 4% PFA for 7 days at room temperature, washed three times with 1X PBS and stored in 70% EtOH at 4°C. The soft tissues were removed manually before analysis.

### Histological staining

Staining was carried out by the Montpellier Experimental Histology Network platform (RHEM) using standardized procedure based on Leica Autostainer XL technology. For each organ sample, 3 µm of thickness slices were produced and mounted on slides. For each animal sampled, Hematoxylin, Eosin and Saffron (HES) staining was performed to assess the of tissue’s structural parameters. Sirius Red (SR) and Masson’s Trichrome (MT) staining were also used to measure the fibrosis level in those tissues. All slides were scanned by the Montpellier Ressources Imagerie platform (MRI), using a Hamamatsu Photonics NanoZoomer, fitted with a dry x40 objective. NDP.view2 software (Hamamatsu) was used for viewing virtual scans for image acquisition. For all the experiments, acquisition and display settings were identical between each sample to guarantee a reliable quantification.

### DNA methylation

Genomic DNA (gDNA) was extracted from animal lysed tissues using Invitrogen TRIzol™ reagent experimental protocol for DNA isolation (Catalog Numbers 15596026) enabling to isolate sequentially RNA, DNA and proteins from the same sample. gDNA was further processed by Life & Brain GmbH Platform Genomics for DNA methylation profiling. Briefly, bisulfite conversion was performed on qualified gDNA and DNA Methylation levels were measured on “Illumina Infinium Mouse Methylation arrays” according to the manufacturer’s instructions. This new array quantitatively targets over 280 000 CpGs sites across the mice genome. Methylation and Gene Ontology (GO) analysis were conducted using Partek® Genomics Suite® software. We used Illumina’s standard normalization with filtering option to exclude probe on sexual chromosomes and low-quality probes based on p-value detection. For each CpG locus, normalized methylation levels (β values), ranging from 0 (completely unmethylated) to 1 (completely methylated) were calculated and used for differential methylation analysis between control (CTL, n=3) and doxycycline (DOX, n=3) mice group.

A one-way-ANOVA statistical test was performed on log2 converted β values (M value = log2(β/(1-β)). The significant differentially methylated CpG loci (p≤0.01) were used in tissues specific hierarchical clustering, flower diagram and GO analysis.

Two successive statistical tests were used to select the 100 loci multi organs signature. One-way ANOVA (p<0.05) CpG was first applied on M value followed by an additional unpaired nonparametric Mann-Whitney statistical test on the corresponding β values. CpG corresponding to the 100 most significant Mann-Whitney p-value were selected (p<0.0005) and CpG represented in hierarchical clustering to illustrate this “all organs’ signature”.

CpG loci β values corresponding to the 3 highest methylation differences between control (CTL, n=18) and doxycycline (DOX, n=18) mice group were selected from “all organs’ signature” and represented in a scatter plot (**** p<0.0001; according to unpaired – Mann-Whitney test, two-tailed).

### Measurement of histopathological parameters

The architecture of the marginal zone was evaluated according to the scoring method developed previously for evaluation of age-related alteration in mice(58). For each animal 5 to 8 whole white pulp follicles were analyzed. For each of them, the percentage of distortion of the interface between the white pulp and the marginal zone was measured and assigned a score of 1 to 4. In addition to this, the percentage of radial distortion, representing the advance of the marginal zone towards the center of the follicle, was also measured and assigned a second score from 1 to 4. These two scores were then added together to obtain the final score characterizing the alteration of the marginal zone of the follicle. The characterization of the alteration ranges from minimal (score between 7-8) to severe (score between 0-2).

To calculate the mesangial area around the renal glomeruli, 10 to 22 glomeruli were analyzed per animal. The measurement was performed using ImageJ software (NIH).

For the thickness measurement of the various skin layers, images with 20x digital magnification were taken on a minimum of 3 identically oriented skin sections per individual. Hundreds of measurements were then carried out for each layer of the skin. All distance measurements were made with ImageJ.

### Tissue fibrosis

Images were analyzed on ImageJ software equipped with the Fiji processing software using an automated analysis script developed within the laboratory. MT Staining was used to quantify pulmonary fibrosis, and the SR staining was used to measure fibrosis in the kidneys, spleen and heart. A color deconvolution was applied according to the following parameters (Colour_1 R: 0.7995107, G: 0.5913521, B: 0.10528667; Colour_2 R: 0.099971585, G: 0.73738605, B: 0.6680326; Cololour_3 R: 0.59227383, G: 0.3264422, B: 0.73669) for each MT image, using the appropriate function available on Fiji. For all RS images the color deconvolution parameters were set as follow (Colour_1 R: 0.148, G: 0.772, B: 0.618; Colour_2 R: 0.462, G: 0.602, B: 0.651; Cololour_3 R: 0.187, G: 0.523, B: 0.831) based on the MRI developed plugin (http://dev.mri.cnrs.fr/projects/imagej-macros/wiki/Fibrosis_Tool). Then channels of interest were selected and the background noise was subtracted using the MaxEnthropy auto-threshold method. The areas covered by fibrosis were then measured and quantified.

Pulmonary fibrosis was quantified in the superior and inferior right lobes and in the left lobe with 20x digital magnification. Cardiac fibrosis was quantified in the left ventricle. Fibrosis in the spleen was measured in white pulp follicles with a 40x digital magnification. Renal fibrosis was measured in the cortex, medulla and papillae with 20x digital magnification.

### Bone structure evaluation

Osteoporosis and surface degradation were measured on the trabecular and subchondral region (lateral and median plateau in the knee joint) of the tibia. The samples were scanned by X-ray microtomography (µ-CT) on a SkyScan 1176 scanner (Bruker) using CTAn software (Bruker) and the following parameters (aluminum filter, 45kV, 500 µA, resolution of 18µm, 0.5° rotation angle). Scans were then reconstructed using NRecon software (Bruker). The 3D images of the joints were reconstructed using Avizo software (Avizo Lite 9.3.0, FEI Visualization Sciences Group).

### Cartilage degradation

A Leica Microsystems TCS SP5-II confocal laser scanning microscope was used to acquire images of the articular cartilage of the lateral and median plateau. The cartilage was scanned in-depth (XYZ mode) using the following parameters (voxel size 6 μm, 5x dry objective and UV laser light source at 405 nm). Image stacks were used to reconstruct a 3D image of the cartilage as well as for quantification.

### Statistical analyses

For the µ-CT and confocal laser scanning microscopy experiments, each sample was independent and represented an experimental unit providing a unique result.

Statistical analyzes were performed with Prism 7 software (GraphPad). In all histograms, data are presented as the mean ± SEM. For the comparisons of two groups, an unpaired (*in vivo*) or paired (*in vitro*) t-test was used. For the comparisons of the survival curves a log-rank test (Mantel-Cox) was used.

## References

1. J. Campisi, Aging, cellular senescence, and cancer. Annu Rev Physiol 75, 685–705 (2013).

2. C. López-Otín, M. A. Blasco, L. Partridge, M. Serrano, G. Kroemer, The hallmarks of aging. Cell 153, 1194–1217 (2013).

3. S Horvath, K. Raj, DNA methylation-based biomarkers and the epigenetic clock theory of ageing. Nat Rev Genet 19, 371–384 (2018).

4. R. E. Marioni et al., DNA methylation age of blood predicts all-cause mortality in later life. Genome Biol 16, 25 (2015).

5. T. Wang et al., Epigenetic aging signatures in mice livers are slowed by dwarfism, calorie restriction and rapamycin treatment. Genome Biol 18, 57 (2017).

6. D. A. Petkovich et al., Using DNA Methylation Profiling to Evaluate Biological Age and Longevity Interventions. Cell Metab 25, 954-960.e956 (2017).

7. T. M. Stubbs et al., Multi-tissue DNA methylation age predictor in mouse. Genome Biol 18, 68 (2017).

8. B. G. Childs et al., Senescent cells: an emerging target for diseases of ageing. Nat Rev Drug Discov 16, 718–735 (2017).

9. K Takahashi, S. Yamanaka, Induction of pluripotent stem cells from mouse embryonic and adult fibroblast cultures by defined factors. Cell 126, 663–676 (2006).

10. D. A. Robinton, G. Q. Daley, The promise of induced pluripotent stem cells in research and therapy. Nature 481, 295–305 (2012).

11. T. J. Sarkar et al., Transient non-integrative expression of nuclear reprogramming factors promotes multifaceted amelioration of aging in human cells. Nat Commun 11, 1545 (2020).

12. A. Ocampo et al., In Vivo Amelioration of Age-Associated Hallmarks by Partial Reprogramming. Cell 167, 1719-1733.e1712 (2016).

13. P Scaffidi, T. Misteli, Lamin A-dependent nuclear defects in human aging. Science 312, 1059–1063 (2006).

14. F. G. Osorio et al., Splicing-directed therapy in a new mouse model of human accelerated aging. Sci Transl Med 3, 106ra107 (2011).

15. B. Liu et al., Genomic instability in laminopathy-based premature aging. Nat Med 11, 780–785 (2005).

16. B. W. Carey, S. Markoulaki, C. Beard, J. Hanna, R. Jaenisch, Single-gene transgenic mouse strains for reprogramming adult somatic cells. Nat Methods 7, 56–59 (2010).

17. M. Abad et al., Reprogramming in vivo produces teratomas and iPS cells with totipotency features. Nature 502, 340–345 (2013).

18. R Martins, G. J. Lithgow, W. Link, Long live FOXO: unraveling the role of FOXO proteins in aging and longevity. Aging Cell 15, 196–207 (2016).

19. A. M. Leidal, B. Levine, J. Debnath, Autophagy and the cell biology of age-related disease. Nat Cell Biol 20, 1338–1348 (2018).

20. U. G. Kyle et al., Age-related differences in fat-free mass, skeletal muscle, body cell mass and fat mass between 18 and 94 years. European journal of clinical nutrition 55, 663–672 (2001).

21. J. N. Justice et al., Battery of behavioral tests in mice that models age-associated changes in human motor function. Age (Dordr) 36, 583–592 (2014).

22. C. C. Zouboulis, E. Makrantonaki, Clinical aspects and molecular diagnostics of skin aging. Clinics in dermatology 29, 3–14 (2011).

23. W. E. Roberts, Skin type classification systems old and new. Dermatologic clinics 27, 529–533, viii (2009).

24. Y Dor, H. Cedar, Principles of DNA methylation and their implications for biology and medicine. Lancet 392, 777–786 (2018).

25. J. A. Viscarra, Y. Wang, H. P. Nguyen, Y. G. Choi, H. S. Sul, Histone demethylase JMJD1C is phosphorylated by mTOR to activate de novo lipogenesis. Nat Commun 11, 796 (2020).

26. S. Watanabe et al., JMJD1C demethylates MDC1 to regulate the RNF8 and BRCA1-mediated chromatin response to DNA breaks. Nat Struct Mol Biol 20, 1425–1433 (2013).

27. M. Tiana et al., The SIN3A histone deacetylase complex is required for a complete transcriptional response to hypoxia. Nucleic acids research 46, 120–133 (2018).

28. V. L. Barnes et al., SIN3 is critical for stress resistance and modulates adult lifespan. Aging (Albany NY) 6, 645–660 (2014).

29. L Zhang, N. Stokes, L. Polak, E. Fuchs, Specific microRNAs are preferentially expressed by skin stem cells to balance self-renewal and early lineage commitment. Cell Stem Cell 8, 294–308 (2011).

30. D. J. Hunter, S. Bierma-Zeinstra, Osteoarthritis. Lancet 393, 1745–1759 (2019).

31. T. J. Aspray, T. R. Hill, Osteoporosis and the Ageing Skeleton. Sub-cellular biochemistry 91, 453–476 (2019).

32. H. L. Stewart, C. E. Kawcak, The Importance of Subchondral Bone in the Pathophysiology of Osteoarthritis. Frontiers in veterinary science 5, 178 (2018).

33. A. Birbrair et al., Type-1 pericytes accumulate after tissue injury and produce collagen in an organ-dependent manner. Stem cell research & therapy 5, 122 (2014).

34. M. S. Espindola et al., Targeting of TAM Receptors Ameliorates Fibrotic Mechanisms in Idiopathic Pulmonary Fibrosis. Am J Respir Crit Care Med 197, 1443–1456 (2018).

35. M. V. Nastase, J. Zeng-Brouwers, M. Wygrecka, L. Schaefer, Targeting renal fibrosis: Mechanisms and drug delivery systems. Adv Drug Deliv Rev 129, 295–307 (2018).

36. A Lemmer, L. B. VanWagner, D. Ganger, Assessment of Advanced Liver Fibrosis and the Risk for Hepatic Decompensation in Patients With Congestive Hepatopathy. Hepatology (Baltimore, Md.) 68, 1633–1641 (2018).

37. L Li, Q. Zhao, W. Kong, Extracellular matrix remodeling and cardiac fibrosis. Matrix biology : journal of the International Society for Matrix Biology 68-69, 490–506 (2018).

38. J. M. van Deursen, The role of senescent cells in ageing. Nature 509, 439–446 (2014).

39. P. K. Naik, B. B. Moore, Viral infection and aging as cofactors for the development of pulmonary fibrosis. Expert review of respiratory medicine 4, 759–771 (2010).

40. P. W. Noble, C. E. Barkauskas, D. Jiang, Pulmonary fibrosis: patterns and perpetrators. J Clin Invest 122, 2756–2762 (2012).

41. L. Palacio et al., Restored immune cell functions upon clearance of senescence in the irradiated splenic environment. Aging Cell 18, e12971 (2019).

42. J. H. Lim et al., Age-associated molecular changes in the kidney in aged mice. Oxidative medicine and cellular longevity 2012, 171383 (2012).

43. A. Rodríguez-Matellán, N. Alcazar, F. Hernández, M. Serrano, J. Ávila, In Vivo Reprogramming Ameliorates Aging Features in Dentate Gyrus Cells and Improves Memory in Mice. Stem Cell Reports 15, 1056–1066 (2020).

44. Y. Lu et al., Reprogramming to recover youthful epigenetic information and restore vision. Nature 588, 124–129 (2020).

45. N Olova, D. J. Simpson, R. E. Marioni, T. Chandra, Partial reprogramming induces a steady decline in epigenetic age before loss of somatic identity. Aging Cell 18, e12877 (2019).

46. K. Ohnishi et al., Premature termination of reprogramming in vivo leads to cancer development through altered epigenetic regulation. Cell 156, 663–677 (2014).

47. E. Senís et al., AAV vector-mediated in vivo reprogramming into pluripotency. Nature Communications 9, 2651 (2018).

48. M. C. Doeser, H. R. Schöler, G. Wu, Reduction of Fibrosis and Scar Formation by Partial Reprogramming In Vivo. Stem Cells 36, 1216–1225 (2018).

49. A. Chiche et al., Injury-Induced Senescence Enables In Vivo Reprogramming in Skeletal Muscle. Cell Stem Cell 20, 407-414.e404 (2017).

50. L. Mosteiro et al., Tissue damage and senescence provide critical signals for cellular reprogramming in vivo. Science 354 (2016).

51. L. Lapasset et al., Rejuvenating senescent and centenarian human cells by reprogramming through the pluripotent state. Genes Dev 25, 2248–2253 (2011).

52. R. de Cabo, D. Carmona-Gutierrez, M. Bernier, M. N. Hall, F. Madeo, The search for antiaging interventions: from elixirs to fasting regimens. Cell 157, 1515–1526 (2014).

53. J. Campisi et al., From discoveries in ageing research to therapeutics for healthy ageing. Nature 571, 183–192 (2019).

54. J. L. Kirkland, T. Tchkonia, Senolytic drugs: from discovery to translation. Journal of internal medicine 288, 518–536 (2020).

55. D. J. Baker et al., Clearance of p16Ink4a-positive senescent cells delays ageing-associated disorders. Nature 479, 232–236 (2011).

56. D. J. Baker et al., Naturally occurring p16(Ink4a)-positive cells shorten healthy lifespan. Nature 530, 184–189 (2016).

57. M. Xu et al., Senolytics improve physical function and increase lifespan in old age. Nat Med 24, 1246–1256 (2018).

58. S. Z. Birjandi, J. A. Ippolito, A. K. Ramadorai, P. L. Witte, Alterations in marginal zone macrophages and marginal zone B cells in old mice. J Immunol 186, 3441–3451 (2011).

59. P. M. Quiros, A. Goyal, P. Jha, J. Auwerx, Analysis of mtDNA/nDNA Ratio in Mice. Current protocols in mouse biology 7, 47–54 (2017).

60. G. Morawietz et al., Revised guides for organ sampling and trimming in rats and mice--Part 3. A joint publication of the RITA and NACAD groups. Experimental and toxicologic pathology : official journal of the Gesellschaft fur Toxikologische Pathologie 55, 433–449 (2004).

